# Zbtb38 transcriptionally activates XIAP to regulate apoptosis in development and cancer

**DOI:** 10.1101/2024.09.05.611337

**Authors:** Toshiaki Shigeoka, Hiroyuki Nagaoka, Nunuk Aries Nurulita, Shogo Tada, Yasumasa Bessho, Yasumasa Ishida, Eishou Matsuda

**Affiliations:** Functional Genomics and Medicine, Nara Institute of Science and Technology, Ikoma, 630-0192, Japan; Parexel International Inc. Tradepia Yodoyabashi, 2-5-8, Imabashi, Chuo-ku, Osaka-shi, Osaka 541-0042, Japan; Faculty of Pharmacy, University of Muhammadiyah Purwokerto, Jalan Raya Dukuhwaluh PO BOX 202, 53052 Purwokerto, Central Java, Indonesia; Gene Regulation Research, Nara Institute of Science and Technology, Ikoma, 630-0192, Japan

**Keywords:** Apoptosis, XIAP, Bcl-2, embryonic stem cell differentiation, embryonic development, and cancer

## Abstract

The X-linked inhibitor of apoptosis protein (XIAP) is a key suppressor of apoptosis, crucial for cellular differentiation, embryogenesis, and cancer progression. However, its upstream regulatory mechanisms remain poorly understood. Here, we demonstrate that the zinc finger transcription factor Zbtb38, a negative regulator of apoptosis, modulates XIAP expression in both loss- and gain-of-function experiments, irrespective of p53 expression. Notably, XIAP overexpression rescues the apoptosis induced by Zbtb38 knockdown, indicating that Zbtb38-associated apoptosis is at least partially XIAP-dependent. Mechanistically, Zbtb38 binds to E-box motifs within upstream regulatory regions of XIAP and activates its transcription. During embryonic stem (ES) cell differentiation and embryogenesis, Zbtb38 depletion increases apoptosis and reduces XIAP and Bcl-2 expression, underscoring their functional relevance in these processes. Analysis of human tumor datasets reveals a strong positive correlation between ZBTB38 and XIAP expression, with elevated ZBTB38 levels associated with high-grade malignancies. Furthermore, Zbtb38 knockdown induces apoptosis in cancer cells with reduced XIAP expression, regardless of p53 expression. Collectively, these findings uncover a novel Zbtb38-XIAP axis that regulates apoptosis during cellular differentiation, development, and oncogenesis, and highlight its therapeutic potential in XIAP-driven and p53-deficient tumors.

## INTRODUCTION

In mammals, apoptosis is a genetically programmed form of cell death essential for maintaining tissue homeostasis by eliminating damaged or unnecessary cells [1] [2]. It also contributes to immune surveillance by removing abnormal or neoplastic cells. Intrinsic apoptosis, also known as mitochondrial apoptosis, is triggered by internal signals such as DNA damage, hypoxia, hydrogen peroxide, and survival factor deprivation [3][4]. These stimuli activate the initiator caspase-9, which is cleaved into its active form (cCasp9) and subsequently activates downstream effector caspases, including caspase-3 and caspase-7 (cCasp3 and cCasp7) [5][6]. This caspase cascade results in the cleavage of various cellular substrates, with cleaved poly (ADP-ribose) polymerase (cPARP) serving as a hallmark of irreversible apoptosis [7].

The Bcl-2 family of proteins plays a central role in regulating the release of apoptogenic proteins from the mitochondria. This family includes anti-apoptotic proteins (e.g., Bcl-2, Bcl-xL, and Bcl-w), BH3-only pro-apoptotic proteins (e.g., Bik, Bad, Noxa, and Puma), and multidomain pro-apoptotic proteins (e.g., Bax and Bak) [8]. BH3-only proteins promote apoptosis by binding to and inhibiting anti-apoptotic proteins [9][10]. Bcl-2, a well-characterized anti-apoptotic protein, sequesters Bax and Bak to prevent cytochrome c release from mitochondria [4][5]. Another key regulator is p53, which promotes apoptosis by transcriptionally activating pro-apoptotic genes (e.g., Noxa and Puma), repressing anti-apoptotic genes (e.g., Bcl-2 and Bcl-xL), and interacting directly with mitochondrial membranes [11]. These multifaceted functions underscore p53’s pivotal role as a tumor suppressor in cancer [12]. In contrast to the Bcl-2 family proteins, the inhibitor of apoptosis proteins (IAPs) suppress apoptosis by directly inhibiting caspases. The IAP family comprises three classes: Class I (cIAP1, cIAP2, and XIAP), Class II (NAIP), and Class III (survivin and Bruce) [13][14]. Class I IAPs contain baculovirus IAP repeat (BIR) domains that directly bind and inhibit caspases. Notably, XIAP uniquely inhibits caspases at physiological concentrations, distinguishing it from cIAP1 and cIAP2 [15][16]. While single knockout (KO) mice for XIAP, cIAP1, or cIAP2, and XIAP/cIAP2 double KOs are viable and fertile, double KO mice for XIAP/cIAP1 or cIAP1/cIAP2 exhibit embryonic lethality around embryonic day 10.5 (E10.5), indicating functional redundancy [17][18][19]. IAP inhibitors such as Smac and Xaf1 displace caspases by binding to IAP BIR domains [20], while Htra2 promotes IAP degradation through its serine protease activity [21]. Despite XIAP’s central role in apoptosis inhibition, its upstream regulatory mechanisms remain poorly defined.

We and others have shown that Zbtb38 (also known as CIBZ) binds to both methylated and unmethylated DNA through its first zinc finger cluster (ZF1–5), functioning as either a transcriptional activator or repressor depending on the chromatin context and target gene [22][23][24][25][26]. Our previous work demonstrated that Zbtb38 promotes the proliferation of mouse ES cells while inhibiting their differentiation toward the mesodermal lineage [27][28]. We also reported that siRNA-mediated knockdown of Zbtb38 induces apoptosis in murine cells, as evidenced by increased levels of cCasp9, cCasp3, cCasp7, and cPARP, along with enhanced annexin V uptake [29]. Consistently, Zbtb38 overexpression suppresses apoptosis and restores spinal cord function after injury in mice [30]. Recently, we showed that heterozygous loss of Zbtb38 results in increased apoptosis shortly after implantation, leading to early embryonic lethality at E9.5 [31]. Despite these findings, the molecular mechanism by which Zbtb38 regulates apoptosis remains unclear.

In this study, we aimed to identify the downstream targets of Zbtb38 during apoptosis and elucidate its regulatory mechanisms. To this end, we performed comparative gene expression analyses in p53-expressing and p53-deficient cells, including cancer cell lines, following Zbtb38 knockdown and overexpression. Since Zbtb38 expression consistently correlated with XIAP levels across various cell lines, suggesting that XIAP as a potential target. Sequence analysis revealed two conserved E-box motifs upstream of the XIAP transcription start site (TSS). Chromatin immunoprecipitation (ChIP) and luciferase assays confirmed Zbtb38 binding to these E-box regions and activation of XIAP transcription. Analysis of human tumor datasets revealed a strong ZBTB38-XIAP expression correlation, with elevated ZBTB38 in high-grade tumors. Moreover, Zbtb38 depletion increased apoptosis and reduced XIAP and Bcl-2 levels during ES cell differentiation and embryogenesis, highlighting their developmental roles. Together, these findings demonstrate that Zbtb38 regulates apoptosis by activating XIAP transcription via E-box motifs during cellular differentiation, development, and tumorigenesis.

## RESULTS

### XIAP identified as a potential direct target of Zbtb38 in apoptosis regulation

Our previous studies demonstrated that siRNA-mediated knockdown of Zbtb38 induces apoptosis in C2C12 cells and p53 KO mouse embryonic fibroblasts (MEFs). However, its direct downstream target remains unclear. To address this, we analyzed the effects of Zbtb38 knockdown in p53-expressing (MEFs, NMuMG, C2C12) and p53-deficient (p53 KO and p53/Dnmt1 double knockout [DKO]) cell lines. Since two independent siRNA duplexes targeting distinct regions of the Zbtb38 gene produced consistent results in preliminary experiments [29], one was used for subsequent experiments. RT-qPCR analysis revealed that Zbtb38 knockdown consistently reduced XIAP mRNA levels across all cell lines, with Bcl-2 expression decreased in NMuMG and C2C12 cells (Fig. 1). In contrast, the expression levels of other IAP genes (cIAP1, cIAP2, survivin), IAP inhibitor genes (Smac, HtrA2, XAF1), and additional apoptosis-related genes exhibited minimal or inconsistent changes (Fig. 1, S1A). For example, Zbtb38 knockdown upregulated Bik in MEFs and Noxa in p53 KO MEFs, while increasing the expression of both anti-apoptotic (Bcl-xL, Bcl-w) and pro-apoptotic genes (Bax, Bak, Bik, Noxa, and Puma) in DKO MEFs (Fig. 1, S1A). These findings suggest that XIAP is a likely direct downstream target of Zbtb38.

**Fig. 1.**
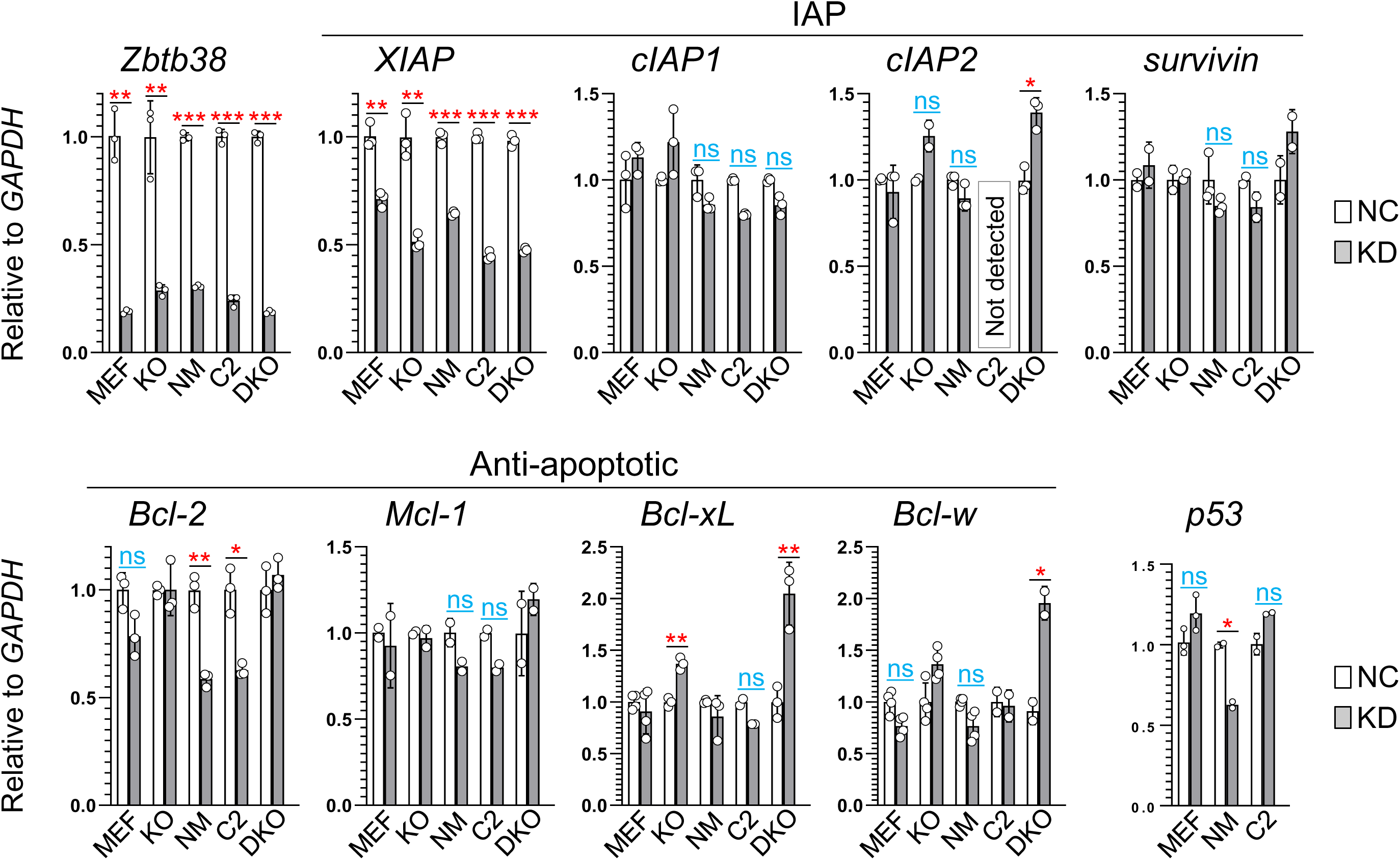
XIAP expression correlates with Zbtb38 expression across all siRNA-mediated knockdown cells. Expression of apoptosis-related genes following Zbtb38 knockdown in p53-expressing and p53-deficient cell lines. mRNA levels were measured by RT-qPCR in negative control (NC, empty bars) and Zbtb38 knockdown (KD, grey bars) groups. GAPDH was used as an internal control, and transcript levels were normalized to GAPDH (set as 1). KO, p53 knockout MEFs; NM, NMuMG; C2, C2C12; DKO, p53/Dnmt1 double knockout MEFs. \**p < 0.05,* \*\**p* < 0.01, \*\*\**p* < 0.001. “ns” indicates no significance.

### Zbtb38 modulates apoptosis through XIAP regulation, independently of p53 expression and DNA methylation

To further investigate whether XIAP is a direct downstream target of Zbtb38, we performed immunoblotting analyses following Zbtb38 knockdown, overexpression, and rescue experiments. Zbtb38 knockdown consistently reduced XIAP protein levels and induced apoptosis, as evidenced by increased levels of cCasp3 and cPARP across all examined cell lines, regardless of p53 expression status (Fig. 2A–C, S1B–C). Notably, Zbtb38 knockdown induced apoptosis in p53/Dnmt1 DKO MEFs, which lack DNA methylation, demonstrating that its anti-apoptotic function is independent of both p53 and DNA methylation (Fig. S1C). To validate XIAP as a functional downstream target, we overexpressed Flag-tagged Zbtb38 (Flag-Zbtb38) in both p53-expression and p53-deficient cells. In C2C12 cells, Flag-Zbtb38 increased XIAP expression and reduced serum deprivation-induced apoptosis compared to controls (Fig. 2D). Similarly, in p53 KO MEFs, Flag-Zbtb38 elevated XIAP expression and inhibited apoptosis triggered by serum deprivation or hydrogen peroxide (H₂O₂) treatment (Fig. 2E–F). Semi-quantitative PCR further confirmed a positive correlation between Zbtb38 and XIAP in both Zbtb38-overexpressing and knockdown C2C12 cells, but not with other apoptosis-related genes (Fig. S2). To test whether XIAP mediates Zbtb38’s anti-apoptotic function, we performed rescue experiments. In C2C12 cells, overexpression of either untagged XIAP or Flag-XIAP significantly suppressed apoptosis resulting from Zbtb38 knockdown (Fig. 2G). In p53 KO MEFs, XIAP overexpression attenuated apoptosis induced by Zbtb38 knockdown in a dose-dependent manner (Fig. 2H). Importantly, XIAP overexpression did not alter Zbtb38 expression, confirming XIAP as a downstream target. Together, these results demonstrate that Zbtb38 suppresses apoptosis primarily through the upregulation of XIAP, independently of p53 expression and DNA methylation.

**Fig. 2.**
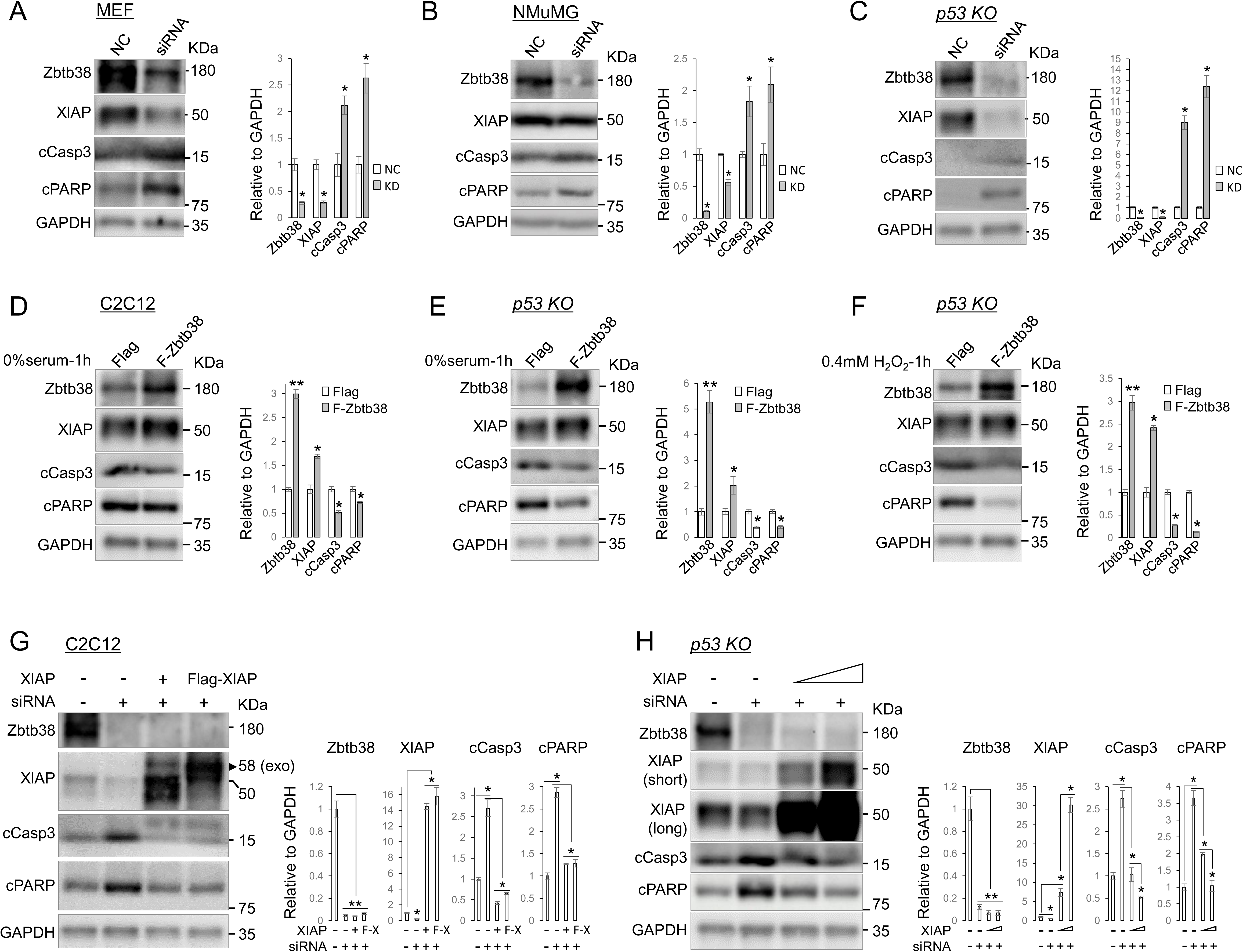
Zbtb38 regulates apoptosis through XIAP, independently of p53 expression and DNA methylation. **A–C** Zbtb38 knockdown induced apoptosis in MEFs **(A)**, NMuMG **(B)**, and p53 KO MEFs **(C)**. NC, negative control (empty bars); KD, Zbtb38 knockdown (grey bars). **D** C2C12 cells were transfected with Flag or Flag-Zbtb38 (F-Zbtb38) for 36 h, followed by serum starvation (0% FBS) for 1 h. **E, F** p53 KO MEFs were transfected with Flag or Flag-Zbtb38 for 30–36 h, followed by serum withdrawal **(E)** or treatment with 0.4 mM H₂O₂ for 1 h **(F)**. **G, H** XIAP overexpression rescued Zbtb38 knockdown-induced apoptosis in C2C12 **(G)** and p53 KO MEFs **(H)**. Cells were transfected with Zbtb38 siRNA or NC for 24 h, followed by XIAP or Flag-XIAP (F-X) transfection for 24 h. Arrows indicate exogenous (exo) XIAP **(G)**. Protein levels of Zbtb38, XIAP, cPARP, and cCasp3 were assessed by immunoblotting. Quantification relative to GAPDH is shown in the right panel of each blot. \**p* < 0.05, \*\*\**p* < 0.001.

### Zbtb38 binds to E-box motifs in the XIAP upstream region and activates transcription

To investigate whether Zbtb38 binds to the XIAP promoter or its upstream regulatory regions, we performed ChIP assays in C2C12 cells. Given the limited understanding of XIAP transcriptional regulation in mammals, we focused on four evolutionarily conserved regions upstream of the TSS, designated as regions a–d (Fig. 3A). ChIP analysis revealed that Zbtb38 specifically bound to regions c and d, located approximately 4.1 kb and 4.7 kb upstream of XIAP’s TSS, respectively, but not to the proximal promoter regions a or b (Fig. 3B). A similar binding pattern was observed in p53 KO MEFs (Fig. S3), indicating that Zbtb38 binding is independent of p53 expression.

**Fig. 3.**
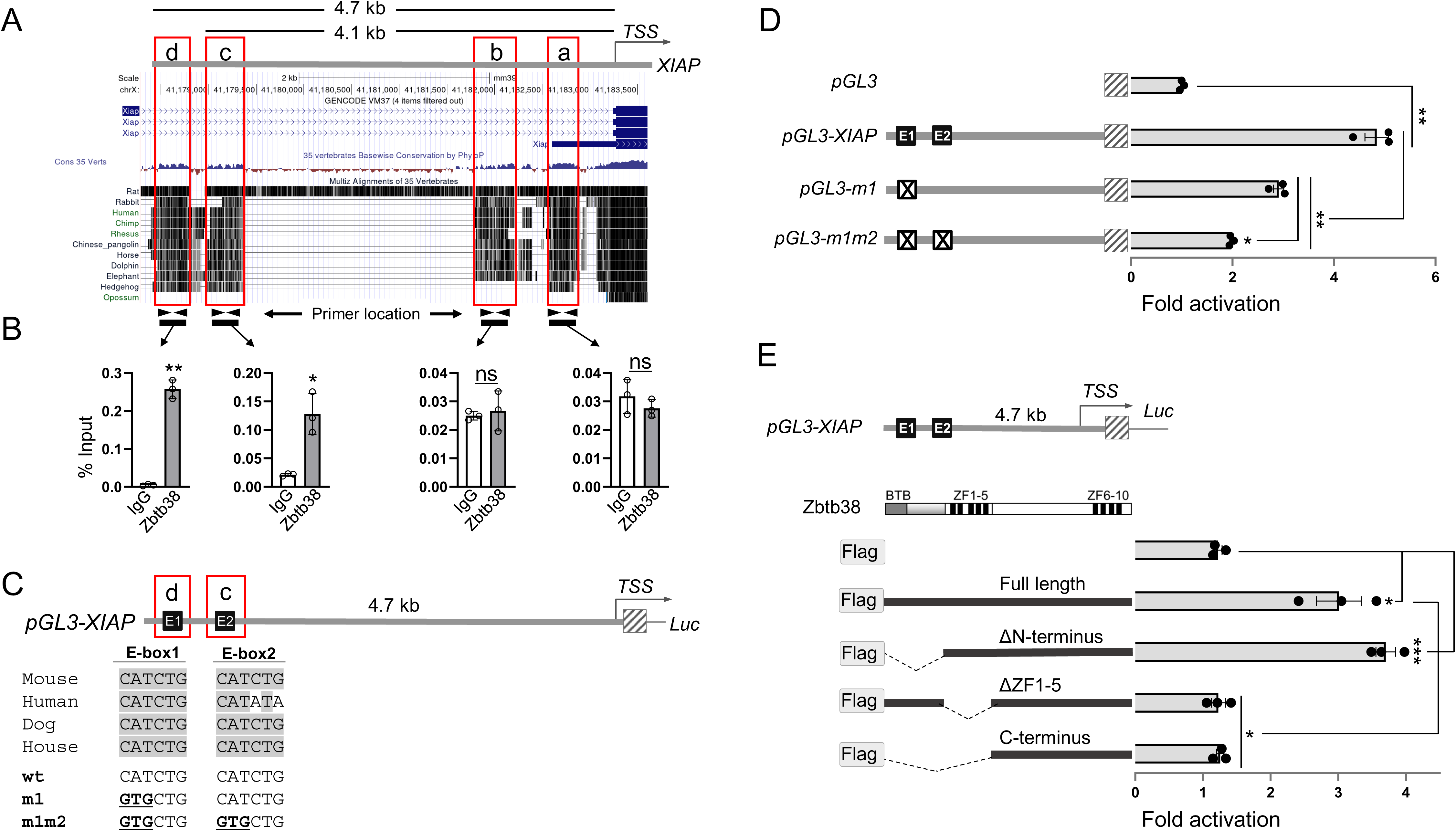
Zbtb38 directly activates XIAP transcription via E-box motifs. **A** Schematic of the XIAP regulatory regions. Red circles indicate conserved regions among vertebrate species. Regions a and b represent putative promoter regions; regions c and d indicate putative enhancer regions. TSS, transcriptional start site. **B** ChIP-qPCR results in C2C12 cells. IgG-precipitated DNA (negative control) and Zbtb38-immunoprecipitated DNA were amplified by RT-qPCR using primers at the indicated locations. Data are shown as fold enrichment relative to the input control (set to 1). **C** Schematic representation of two conserved E-box motifs in the XIAP upstream region. wt, wild-type; m1, E-box1 mutant; m1m2, double mutant of E-box1 and E-box2. Underlined sequences show nucleotide substitutions. Luc, luciferase reporter. **D** Luciferase assay in HEK293T cells co-transfected with pGL3 or indicated pGL3 constructs and Flag or Flag-Zbtb38. Luciferase activity was normalized to that of pGL3/Flag (set to 1). **E** Schematic of Zbtb38 deletion mutants fused to Flag and their transcriptional activation of the pGL3-XIAP 4.7 kb region. Luciferase activity was normalized as in (D). \**p* < 0.05, \*\**p* < 0.01, \*\*\**p* < 0.001. “ns” indicates no significance.

To assess the transcriptional activity of Zbtb38 on XIAP, we performed luciferase assays in HEK293T and C2C12 cells, which express low and high levels of Zbtb38, respectively [25]. Zbtb38 overexpression in HEK293T cells increased luciferase activity of the XIAP 4.7 kb upstream region in a dose-dependent manner, while its knockdown in C2C12 cells suppressed the activity of this region (Fig. S4). Motif analysis identified two canonical E-box motifs (CANNTG), designated E-box1 (E1) and E-box2 (E2), located within regions c and d, respectively (Fig. 3C). Luciferase assays using reporter constructs with point mutations in E1 (m1) or in both E1 and E2 (m1/m2) showed that Zbtb38 strongly activated the wild-type 4.7 kb upstream region, whereas activation was reduced in the m1 mutant and further attenuated in the m1/m2 double mutant (Fig. 3D). These results indicate that both E-boxes contribute to Zbtb38-mediated activation of XIAP, although residual activity in the m1/m2 mutant suggests the involvement of additional regulatory elements. To identify the Zbtb38 domains critical for XIAP activation, we examined a series of deletion mutants. Luciferase assays showed that both full-length Zbtb38 and the ΔN-terminus (containing ZF1–5 but lacking the BTB domain) activated the 4.7 kb region of XIAP, whereas the ΔZF1–5 mutant (lacking ZF1–5) and the C-terminus (lacking both the BTB domain and ZF1–5) nearly lost this activity (Fig. 3E). These results demonstrate that ZF1–5 is essential for XIAP transcriptional activation and that ZF1–5, together with the C-terminus, is sufficient for this function. Supporting this conclusion, immunoblotting showed that full-length Zbtb38 upregulated XIAP expression and suppressed apoptosis induced by serum deprivation, whereas the ΔZF1–5 mutant, despite being expressed at higher levels, failed to do so (Fig. S5). Collectively, these findings confirm that ZF1–5 is critical for Zbtb38-mediated XIAP activation and its anti-apoptotic function.

### Loss of Zbtb38 downregulates XIAP and Bcl-2, leading to increased apoptosis during ES cell differentiation and embryogenesis

We previously reported that Zbtb38 depletion (via knockdown or knockout) in undifferentiated ES cells does not induce apoptosis but instead reduces proliferation, suggesting increased apoptotic susceptibility upon differentiation [27]. To test this hypothesis, we induced differentiation of ES cells in vitro using embryoid body formation (Fig. 4A). RT-qPCR analysis confirmed successful differentiation, as evidenced by the upregulation of ectodermal (Sox2, Nestin), endodermal (Gata4, Gata6), and mesodermal (Brachyury, Mesp1) markers from day 0 (undifferentiated) to day 5 (Fig. S6). Immunoblotting showed progressive reductions in XIAP (days 0–2) and Bcl-2 (days 0–4) protein levels in Zbtb38-deleted cells, coinciding with increased apoptosis from day 2 to 5 (Fig. 4B–C). RT-qPCR revealed decreased XIAP mRNA levels at days 0–2 and reduced Bcl-2 mRNA levels at day 0 in Zbtb38 KO ES cells compared with controls (Fig. 4D). Additionally, expression of cIAP1, cIAP2, and Bcl-w was also reduced in undifferentiated Zbtb38 KO ES cells, whereas the expression of other apoptosis-related genes remained largely unchanged (Fig. 4D, S7). These results suggest that Zbtb38 loss sensitizes ES cells to apoptosis during differentiation by downregulating XIAP and Bcl-2.

**Fig. 4.**
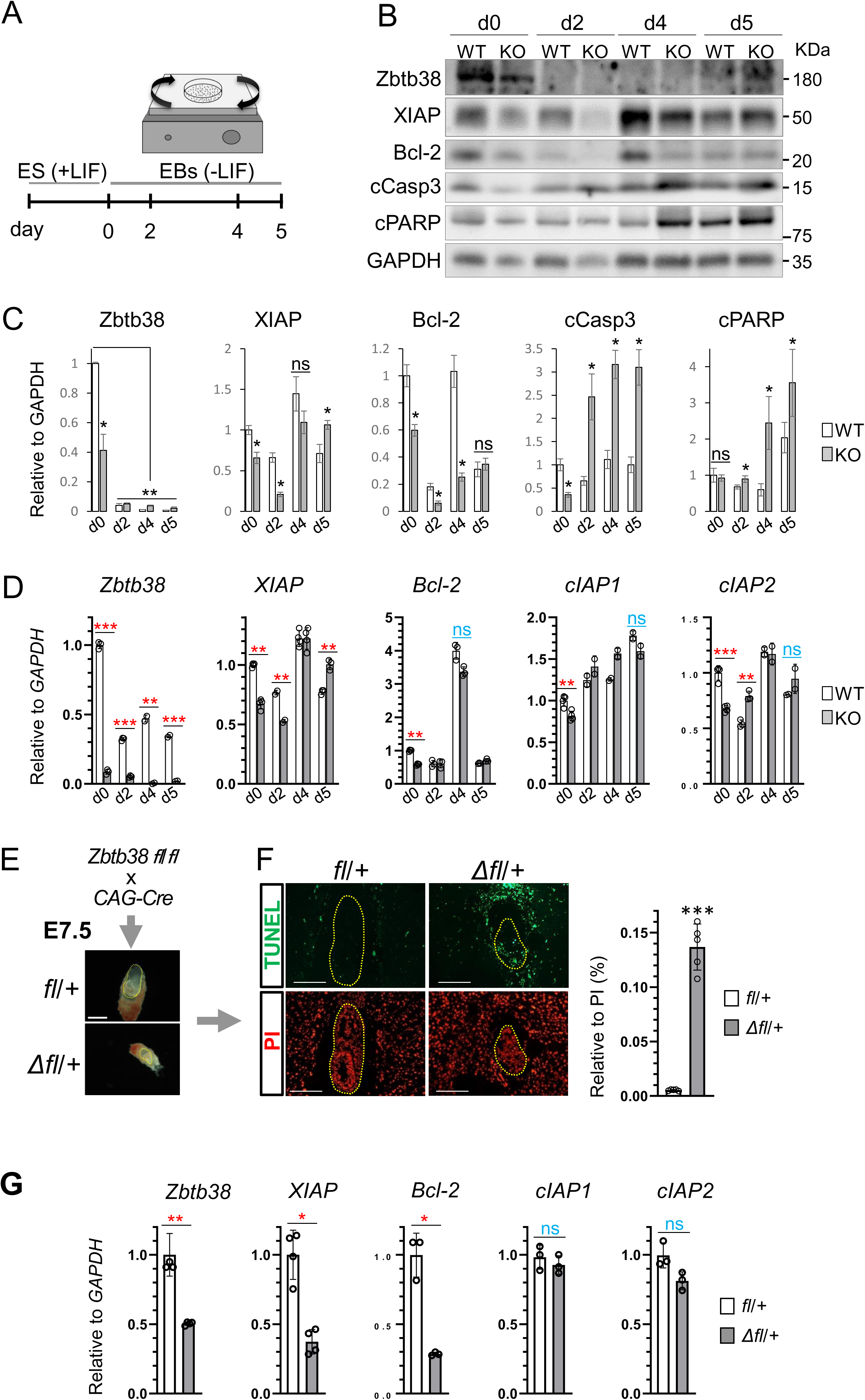
Zbtb38 loss downregulates XIAP and Bcl-2, increasing apoptosis during ES cell differentiation and embryogenesis. **A** Schematic of the ES cell differentiation assay. ES cells were cultured in ES medium (undifferentiated, d0), then differentiated into embryoid bodies (EBs) under suspension culture without LIF. **B, C** Immunoblotting of proteins in wild-type (WT) and Zbtb38 knockout (KO) ES cells at the indicated differentiation days. Protein levels **(B)** were quantified relative to GAPDH and normalized to WT at d0 **(C)**. WT, empty bars; KO, grey bars. **D** RT-qPCR of gene expression in WT and Zbtb38 KO ES cells during differentiation. Transcript levels were normalized to GAPDH. **E** Schematic of the strategy for generating fl/+ and Δfl/+ embryos, with representative images of E7.5 embryos. fl, floxed allele. Scale bar: 100 μm. **F** TUNEL assay of paraffin-embedded E7.5 embryo sections. TUNEL-positive cells (green); nuclei (PI, red). Quantification of TUNEL-positive cells relative to PI-positive nuclei is shown at right. fl/+ (n = 5), Δfl/+ (n = 5). **G** RT-qPCR analysis of Zbtb38 and apoptosis markers in E7.5 embryos. Transcript levels were normalized to GAPDH. fl/+ (n = 3), Δfl/+ (n = 3). \**p* < 0.05, \*\**p* < 0.01, \*\*\**p* < 0.001. “ns” indicates no significant.

To investigate whether this mechanism functions in vivo, we focused on E7.5 embryos because our previous data indicated that increased apoptosis in Zbtb38 heterozygous mutants at this stage [31]. Paraffin-embedded sections from Δfl/+ embryos, generated by intercrossing Zbtb38 fl/fl with CAG-Cre mice, were analyzed using the TUNEL assay to detect DNA fragmentation, a hallmark of apoptosis (Fig. 4E). The results showed that Zbtb38 Δfl/+ embryos exhibited a significantly higher number of TUNEL-positive nuclei compared to controls, correlating with their reduced size (Fig. 4F). RT-qPCR further revealed decreased XIAP and Bcl-2 mRNA levels in Δfl/+ embryos, while cIAP1 and cIAP2 expression remained unchanged (Fig. 4G). Collectively, these data suggest that Zbtb38 loss promotes apoptosis during embryogenesis by downregulating XIAP and Bcl-2.

### Zbtb38 is a potential oncogenic driver and its knockdown triggers apoptosis in cancer cells

Our results demonstrate that Zbtb38 directly suppresses apoptosis via activating XIAP transcription, suggesting a potential role for Zbtb38 in cellular transformation and cancer progression. To further clarify this role, we analyzed gene expression data from The Cancer Genome Atlas (TCGA). Across all TCGA tumor types, ZBTB38 expression exhibited a strong positive correlation with XIAP expression (Fig. 5A). To further explore this relationship at the level of individual tumor samples, we performed t-SNE visualization using transcriptome data from all TCGA samples. Although samples clearly clustered by cancer type (Fig. 5B), expression levels of XIAP and ZBTB38 remained consistently positively correlated within each cancer-specific cluster, indicating that their co-expression is preserved across diverse tumor lineages (Fig. 5C–D). Among the cancer types analyzed, particularly strong correlations were observed in cutaneous melanoma, thyroid carcinoma, prostate adenocarcinoma, breast carcinoma, and colon adenocarcinoma (Fig. 5E). These trends were further validated using independent microarray datasets from the MERAV database (Fig. S8). A genome-wide correlation analysis revealed that ZBTB38 expression was negatively correlated with pro-apoptotic genes such as Bax and BAD, while showing positive correlations with anti-apoptotic genes, with XIAP displaying the strongest correlation (Fig. 5F). Together, these findings indicate that ZBTB38 functions as a negative regulator of apoptosis, thereby promoting cancer cell survival and contributing to malignancy. To better understand the role of ZBTB38 in cancer progression, we also examined its expression across tumor grades. Higher ZBTB38 expression was significantly associated with higher tumor grade (Fig. 5G). Moreover, copy number variation (CNV) analysis revealed that ZBTB38 copy number gain occurred more frequently in high-grade tumors (G3/G4) than in the background CNV distribution (Fig. 5H). These results support ZBTB38 as a potential oncogenic factor and a promising therapeutic target for promoting apoptosis in cancer cells.

**Fig. 5.**
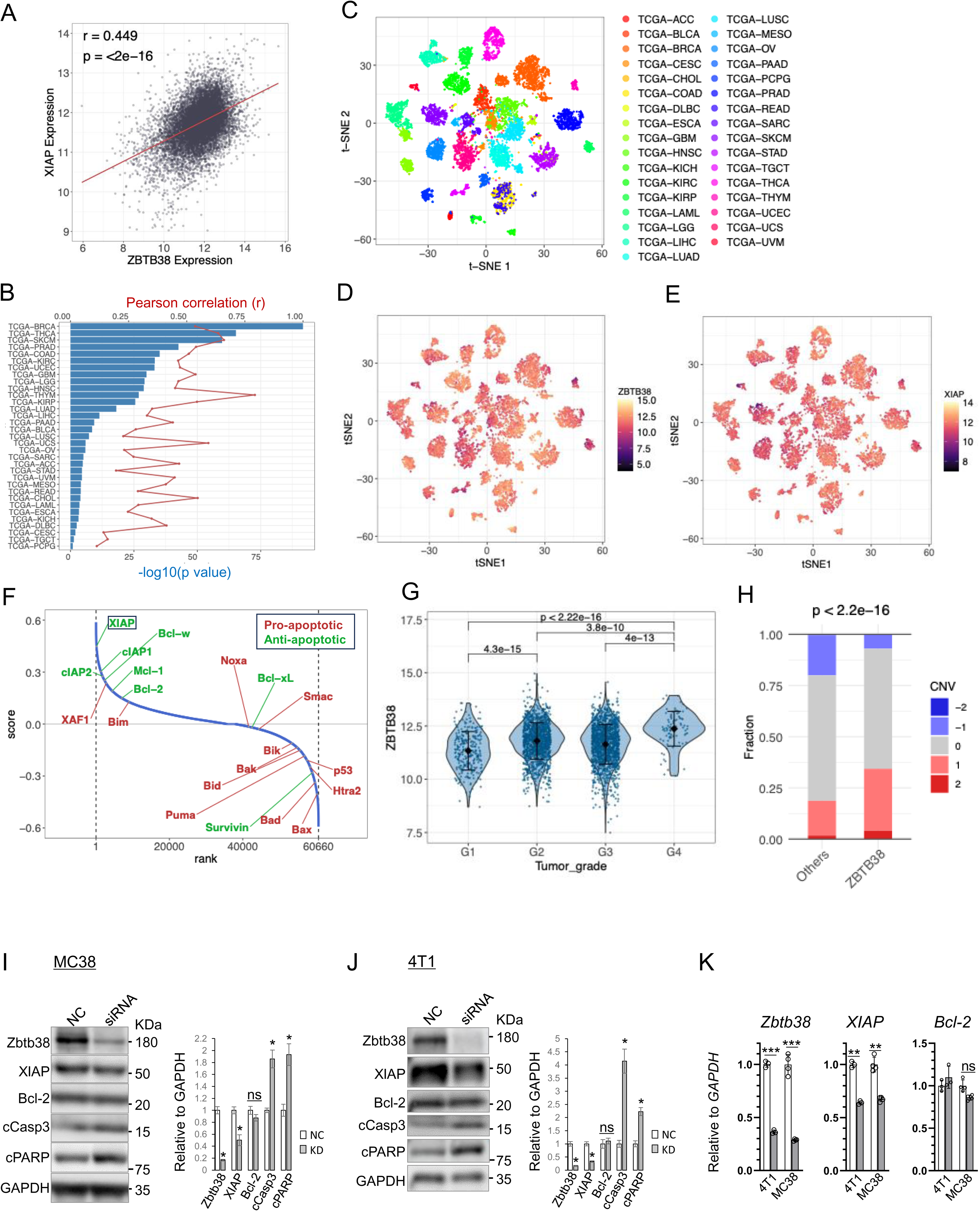
ZBTB38 is a potential oncogenic driver, and its knockdown induces apoptosis in cancer cells. **A** Scatter plot showing correlation between ZBTB38 and XIAP expression across TCGA tumor types. Pearson’s correlation coefficient (r) and p-value are indicated. **B** Correlation between XIAP and ZBTB38 expression across TCGA tumor types (bars: Pearson correlation coefficients; line: -log_10_(*p*-value)). **C** t-SNE projection of transcriptomic data from all TCGA tumor types. **D, E** Expression of ZBTB38 (**D**) and XIAP (**E**) on t-SNE plots. **F** Gene ranking based on co-expression correlation with ZBTB38. Pro- and anti-apoptotic genes are labeled. **G** Distribution of ZBTB38 mRNA expression levels across tumor grades (G1 to G4) in all TCGA tumor types. **H** Fractions of copy number variation (CNV) in high-grade tumors (G3 and G4) for all genes and for ZBTB38. *P*-value was calculated using a Chi-square test (df = 4). **I, J** siRNA-mediated knockdown of Zbtb38 in MC38 and 4T1 cells. Expression of the indicated proteins was assessed by immunoblotting. GAPDH was used as a loading control, and quantification relative to GAPDH is shown in the right panel of each blot. NC, empty bars; KD, grey bars. **K** RT-qPCR analysis of gene expression in 4T1 and MC38 cells. WT (empty column) and KO (Zbtb38 knockout, gray column) cells were used. Each transcript was normalized to GAPDH, set to 1. \**p* < 0.05, \*\**p* < 0.01, \*\*\**p* < 0.001. “ns” indicates no significance.

To directly assess whether Zbtb38 loss induces apoptosis in cancer cells, we performed knockdown experiments in both p53-proficient MC38 (murine colon adenocarcinoma) and p53-deficient 4T1 (murine stage IV mammary carcinoma) cell lines. Immunoblotting revealed that Zbtb38 knockdown led to decreased XIAP protein levels and increased levels of apoptotic markers in both cell lines, while Bcl-2 levels remained largely unchanged (Fig. 5I–J). RT-qPCR confirmed decreased XIAP mRNA expression following Zbtb38 knockdown, with increased Bak expression, while Bcl-2, p53, and other apoptosis-related genes remained minimally affected (Fig. 5K, S9). These results suggest that Zbtb38 knockdown in cancer cells induces apoptosis primarily through XIAP downregulation, independent of Bcl-2 or p53 expression status.

In conclusion, this study demonstrates that Zbtb38 regulates apoptosis by transcriptionally activating XIAP via E-box motifs in its upstream region, independent of p53 and DNA methylation, across various cell types, including cancer cells. During ES cell differentiation and embryogenesis, loss of Zbtb38 increases apoptotic sensitivity by downregulating both XIAP and Bcl-2 (Fig. 6). Furthermore, ZBTB38 expression correlates strongly with XIAP levels and is elevated in high-grade malignancies. These findings deepen the mechanistic understanding of apoptotic regulation and highlight Zbtb38 as a potential therapeutic target in cancers with dysregulated apoptosis.

**Fig. 6.**
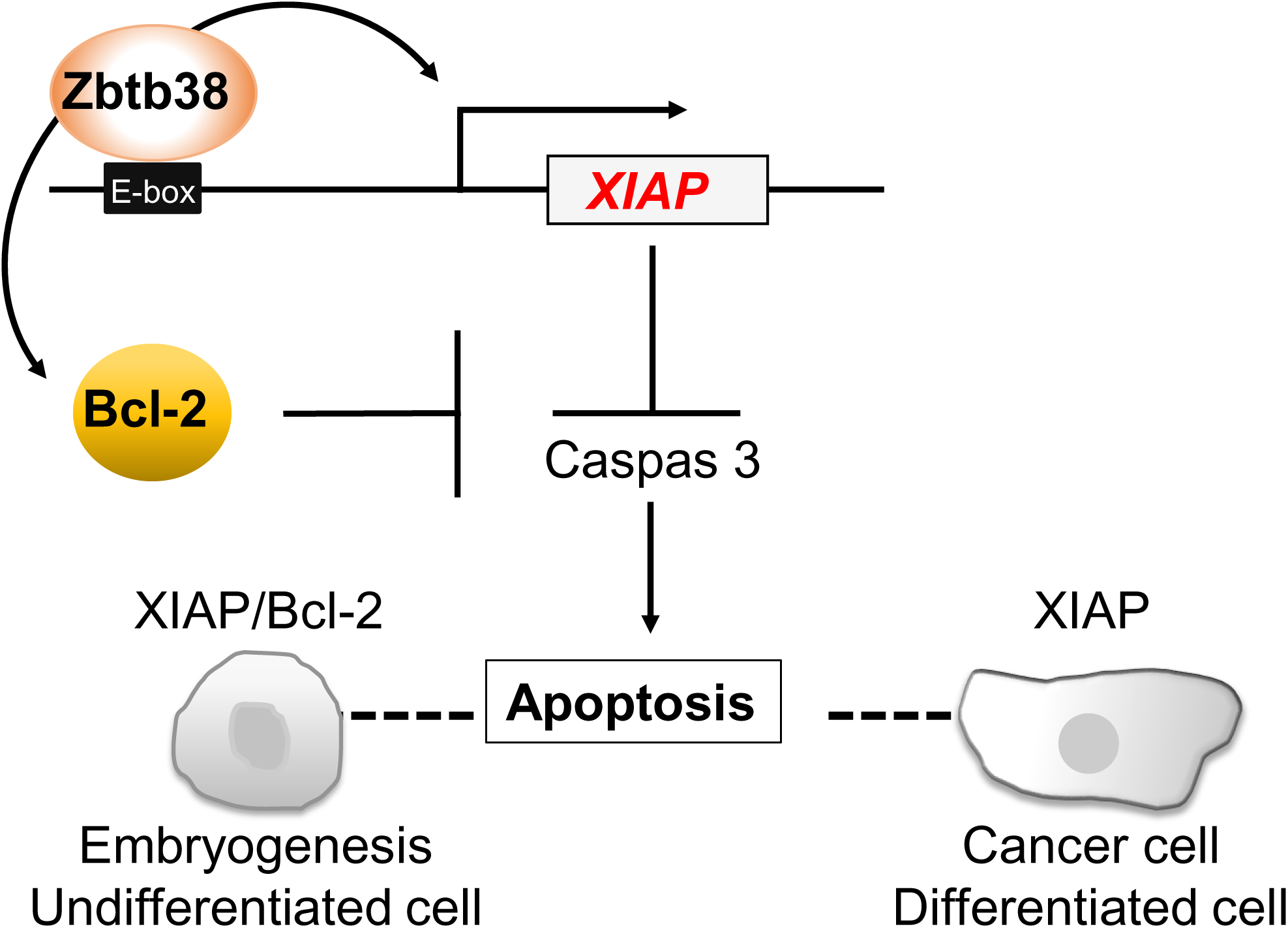
Schematic model of the Zbtb38-XIAP and Zbtb38-XIAP/Bcl-2 axis in apoptosis regulation. Zbtb38 activates XIAP transcription by binding to E-box motifs within the upstream regulatory region of the XIAP gene, thereby inhibiting the activation of caspases and suppressing apoptosis in cancer cells and differentiated cells. During embryogenesis and in undifferentiated cells, both XIAP and Bcl-2—transcriptional targets of Zbtb38—cooperatively inhibit caspase-mediated apoptosis and promote cell survival during development.

## DISCUSSION

This study identifies Zbtb38 as a novel positive regulator of XIAP that functions independently of p53 expression and DNA methylation. The observed correlation between Zbtb38 and XIAP expression across various cell lines, along with the ability of XIAP overexpression to rescue apoptosis in Zbtb38-knockdown cells, establishes XIAP as a key downstream target. These findings are consistent with XIAP’s established role in caspase inhibition and apoptosis suppression, particularly in contexts where cIAP1 and cIAP2 exhibit functional redundancy. Zbtb38’s selective regulation of XIAP, but not other IAPs or IAP inhibitors, underscores its unique role in apoptotic regulation.

The anti-apoptotic function of Zbtb38 appears to be cell type-dependent. In ES cells, loss of Zbtb38 induces apoptosis during differentiation, accompanied by downregulation of XIAP and Bcl-2, suggesting that both proteins are essential for maintaining cell survival during lineage commitment. The temporal sequence, characterized by XIAP and Bcl-2 downregulation preceding apoptosis, supports a “priming” model in which undifferentiated ES cells tolerate Zbtb38 loss but become sensitized to apoptosis upon differentiation. This model is further supported by observations in E7.5 Zbtb38 Δfl/+ embryos, which exhibit increased apoptosis and reduced XIAP and Bcl-2 expression. In contrast, in MEFs and cancer cell lines (e.g., 4T1 and MC38), Bcl-2 levels remain unchanged following Zbtb38 knockdown, suggesting that Bcl-2 is dispensable in these contexts, possibly due to intrinsic cell-type differences or oncogenic reprogramming. Additionally, the differential regulation of pro-apoptotic genes such as Bik and Noxa in p53-deficient cells suggests that Zbtb38 also regulates apoptosis via additional pathways, requiring further investigation.

Mechanistically, ChIP and luciferase assays demonstrate that Zbtb38 binds to E-box motifs within enhancer regions upstream of the XIAP gene and activates transcription via its ZF1–5 domain. This is further supported by an electrophoretic mobility shift assay (EMSA) data showing that ZENON, the rat ortholog of Zbtb38, binds E-box motif through the ZF1–5 domain, which shares 97% sequence identity with that of Zbtb38 [32]. Although Zbtb38 binds to E-box motifs located approximately 4.1–4.7 kb upstream of the XIAP TSS, clear evidence of typical enhancer marks such as p300 binding and H3K27ac enrichment in this region was not observed in publicly available ChIP-seq datasets from ES cells and other cell types. This absence suggests that activation of this region may be context-dependent or transiently induced during specific developmental stages or stress conditions that promote apoptosis. Interestingly, an E-box motif has also been identified in the Bcl-2 promoter, although whether Zbtb38 regulates Bcl-2 through this element remains unclear and requires further study [33]. E-box motifs are canonical binding sites for basic helix-loop-helix (bHLH) transcription factors and are critical for regulating gene expressions in diverse biological contexts, including development, stress responses, and disease [34]. Zbtb38’s ability to activate transcription via E-box motifs expands its functional repertoire beyond methyl-CpG binding, distinguishing it from close homologs such as Zbtb33 (Kaiso) and Zbtb4, which bind methylated DNA or non-methylated Kaiso-binding sequences but not E-box motif [35] [36]. Disrupting the Zbtb38–E-box interaction may therefore offer a novel therapeutic approach to suppress XIAP and promote apoptosis in tumors resistant to current XIAP-targeted therapies. Furthermore, genome-wide association studies (GWAS) have linked ZBTB38 polymorphisms (e.g., rs6763931, which correlates with its expression in blood) to a variety of physiological and pathological traits, including human height, cancer susceptibility, and neurodegenerative disorders [37][38][39][40][41]. These associations highlight the broad physiological and pathological significance of Zbtb38-mediated gene regulation, potentially through its ability to bind to E-box motifs.

XIAP is frequently overexpressed in various cancers and is associated with tumor progression, poor prognosis, and apoptosis resistance [42]. Therapeutic approaches targeting XIAP, such as small-molecule inhibitors and antisense oligonucleotides, have shown promise in preclinical studies [43]. In contrast, the role of Zbtb38 in cancer remains much less defined and appears to be context-dependent. For example, ZBTB38 acts as a tumor suppressor in prostate cancer, where its loss correlates with tumor progression, but functions as an oncogene in bladder cancer, promoting migration, invasion, and survival [44] [45]. In this study, TCGA analysis reveals a strong correlation between ZBTB38 and XIAP expression across multiple tumor types, with ZBTB38 expression elevated in high-grade malignancies. Consistently, Zbtb38 knockdown induced apoptosis in both p53-proficient (MC38) and p53-deficient (4T1) cancer cells, leading to reduced XIAP expression, while Bcl-2 levels remained unchanged. These findings support the conclusion that Zbtb38 mediates apoptosis primarily through XIAP regulation, independently of p53. However, validation in additional cancer cell lines from diverse tumor types is necessary to confirm the generalizability of these findings. This is particularly important given that p53 is mutated or inactivated in approximately 50% of human cancers, contributing to therapeutic resistance. Therefore, targeting the Zbtb38–XIAP axis may offer a promising therapeutic strategy, particularly for p53-deficient tumors where p53-dependent apoptotic pathways are compromised.

In summary, this study demonstrates that Zbtb38 regulates apoptosis by modulating XIAP and, in some contexts, Bcl-2 in a cell type- and context-dependent manner. We show that this function is independent of p53 and DNA methylation, providing new insights into the roles of Zbtb38 in both differentiated and undifferentiated cells, including cancer cells. These findings deepen our mechanistic understanding of apoptotic regulation and highlight Zbtb38 as a potential therapeutic target in XIAP-driven and p53-deficient cancers. Future studies exploring Zbtb38’s broader transcriptional network and context-specific interactions will be essential to further elucidate its physiological and pathological functions.

## MATERIALS AND METHODS

### Generation of Zbtb38 heterozygous (Δfl/+) mice

Zbtb38 Δflox/+ (Δfl/+) mice were generated by randomly crossing six- to eight-week-old female Zbtb38 fl/fl mice with eight- to sixteen-week-old CAG-Cre mice, and vice versa. Embryos of both sexes were used due to the breeding scheme; no sex-specific effects were observed, and data were pooled. Genotyping was performed as previously described [31]. CAG-Cre mice were kindly provided by Dr. Masaru Okabe (Osaka University, Osaka, Japan). Wild-type C57BL/6J mice were obtained from CLEA (Tokyo, Japan). All animal experiments were approved by the Nara Institute of Science and Technology Animal Care Committee and were conducted in accordance with Japanese guidelines.

### Cell culture and ES cell differentiation assay

C2C12 cells were cultured in Dulbecco’s modified Eagle’s medium (DMEM; Nacalai Tesque) supplemented with 15% fetal bovine serum (FBS; Sigma-Aldrich), 2 mM L-glutamine (Nacalai Tesque), and 1% penicillin/streptomycin (Nacalai Tesque) as previously described [25]. p53 KO and p53/Dnmt1 DKO mouse MEFs were kindly provided by Dr. Richard Meehan (MRC Human Genetics Unit, Edinburgh, UK). Immortalized MEFs were established in-house by stable transfection of primary C57BL/6 MEFs with SV40 large T antigen. MEFs, p53 KO MEFs, and p53/Dnmt1 DKO MEFs were maintained in the same medium, supplemented with 10% FBS and 1 mM sodium pyruvate (Invitrogen). NMuMG, 4T1, and HEK293T cells were cultured in DMEM with 10% FBS, 2 mM L-glutamine, and 1% penicillin/streptomycin. MC38 cells were maintained in the same medium with the addition of 100 μM non-essential amino acids and 1 mM sodium pyruvate.

Mouse RF8 (wild-type) and Zbtb38 KO ES cells were cultured as previously described [27]. Briefly, undifferentiated ES cells were grown on mitomycin C-treated SNL-STO feeder cells in DMEM containing 15% FBS, 2 mM L-glutamine, 100 μM non-essential amino acids, 1% penicillin/streptomycin, 0.1 μM β-mercaptoethanol, 1000 IU/ml leukemia inhibitory factor (LIF), and 3i inhibitors (3 μM CHIR99021, 1 μM PD0325901, and 10 μM SB431542; Selleck Chemicals). Differentiation was induced by culturing cells in the same medium without LIF and 3i, allowing embryoid body formation in suspension, as previously described [28]. Mouse ES cells were tested for mycoplasma contamination prior to use, and all cultured cells exhibited normal growth and morphology.

### Reporter constructs and luciferase assays

To generate the pGL3-XIAP 4.7 kb construct, genomic DNA from C2C12 cells was PCR-amplified using primers containing *Kpn*I and *Xho*I sites (Supplementary Table), and the fragment was cloned into the similarly digested pGL3 vector. E-box mutant constructs (pGL3-XIAP m1 and pGL3-XIAP m1/m2) were generated by site-directed mutagenesis (Takara Bio) following the manufacturer’s protocol. All constructs were sequence-verified using an ABI PRISM 3130 genetic analyzer. Primer sequences are listed in the Supplementary Table. Plasmids (pcDNA3-Flag, pcDNA3-Flag-Zbtb38, pcDNA3-Myc, pcDNA3-Myc-Zbtb38, and pcDNA3-Myc-Zbtb38 ΔZF1–5) were prepared as previously described [22]. Luciferase assays were performed as described previously [28]. HEK293T cells were seeded in 24-well plates and transfected with 0–250 ng of expression plasmids and 100 ng of firefly luciferase reporter (pGL3 or pGL3-XIAP 4.7 kb), along with 4 ng of pRL-TK (Renilla luciferase) as an internal control. Total DNA amounts were adjusted with empty pcDNA3. Luciferase activity was measured 32–48 h post-transfection using the Dual-Luciferase Reporter Assay System (Promega), and firefly luciferase values were normalized to Renilla activity. Assays were conducted in duplicate across three independent experiments.

### siRNA and Transient Transfection

Cells were transfected with 5–10 nM Zbtb38-specific Dicer substrate siRNA duplex or a scrambled negative control (Integrated DNA Technologies) using RNAiMAX (Thermo Fisher Scientific), as previously described [29]. Assays including RT-PCR, RT-qPCR, and immunoblotting were conducted 24–48 h post-transfection. Plasmid transfection was performed using Lipofectamine 2000 (Invitrogen), PEI-Max (Polysciences), or ViaFect (Promega), according to the manufacturers’ protocols. Cells were harvested 24–48 h post-transfection.

### RNA Extraction, RT-qPCR, and Semiquantitative RT-PCR

Total RNA was extracted using Sepasol RNA Super G (Nacalai Tesque) for cultured cells or the PicoPure RNA Isolation Kit (Thermo Fisher Scientific) for E7.5 embryos. cDNA synthesis was performed using 500 ng (cultured cells) or ∼5 ng (embryos) of total RNA and ReverTra Ace qPCR RT Master Mix with gDNA Remover (TOYOBO). RT-qPCR was performed using Thunderbird SYBR Green PCR Mix (TOYOBO) on a LightCycler® 96 (Roche), with GAPDH as the internal control. Data were analyzed using the 2-delta-delta Ct method, as previously described [28]. For semiquantitative RT-PCR, reactions were run at 58 °C annealing temperature, with GAPDH as the internal control. PCR products were separated on 2% agarose gels and visualized by ethidium bromide staining. Primer sequences are provided in the Supplementary Table.

### Western Blotting analysis

Protein lysates were prepared in RIPA buffer with protease inhibitors (Roche Diagnostics) as previously described [22]. Equal amounts of protein were separated by 6–15% SDS-PAGE and transferred to PVDF membranes. Blots were probed with antibodies against Zbtb38 [22], XIAP (MBL, M044-3), Bcl-2 (MBL, D038-3), cleaved caspase-3 (cCasp3; Cell Signaling, #9664), cleaved PARP (cPARP; Cell Signaling, #9542), and GAPDH (Proteintech, 60004-1-Ig). Expected molecular weights: Zbtb38 (175 kDa), XIAP (55 kDa), Bcl-2 (26 kDa), cCasp3 (17 kDa), cPARP (89 kDa), GAPDH (36 kDa). HRP-conjugated secondary antibodies (anti-mouse #7076 and anti-rabbit #7074) were from Cell Signaling Technology. Signals were detected using a FUSION chemiluminescence imaging system (Thermo Fisher Scientific).

### ChIP assay and ChIP-qPCR

ChIP assays were performed as previously described [25][28]. Briefly, C2C12 or p53 KO MEFs were cross-linked with 1% formaldehyde, quenched with 0.125 M glycine for 15–30 min. After washing with PBS, cells were lysed in SDS lysis buffer and sonicated on ice to obtain ∼500 bp DNA fragments. Lysates were pre-cleared with Protein G Sepharose beads and immunoprecipitated with anti-Zbtb38 antibody or normal rabbit IgG (negative control). Input DNA (2%) and immunoprecipitated DNA were reverse-crosslinked and analyzed by RT-qPCR using primers listed in the Supplementary Table. ChIP signals were normalized to input DNA. All experiments were performed in triplicate.

### Embryo dissection and TUNEL assay

Morphological and histological analyses of embryos were performed on E7.5 embryos. Bright-field images were captured using a Nikon E800M inverted microscope. For histological analysis, embryos were rinsed with ice-cold PBS, fixed in 4% paraformaldehyde at room temperature for 10-20 min, dehydrated through graded alcohols, and embedded in paraffin. Paraffin sections were used for Terminal deoxynucleotidyl transferase (TdT)-mediated dUTP nick-end labeling (TUNEL) assay (#8445, MBL) as previously described [31]. Briefly, sections were deparaffinized, labeled with TdT and nucleotide mix, counterstained with 1 μg/ml propidium iodide (PI), and mounted. Fluorescent images were acquired using a TCS SP8 confocal microscope (Leica Microsystems).

### Data acquisition and processing for human tumor data analysis

The transcriptome data of human tumors were downloaded from The Cancer Genome Atlas (TCGA) using the TCGA biolinks package (v2.30.4) in R (v4.3.2). Raw count data were processed with DESeq2 (v1.42.1), and a variance stabilizing transformation (VST) was applied prior to downstream analyses. For correlation analyses, Pearson’s correlation coefficients (r) and p-values were calculated in R to assess the association between target gene expression levels. Principal component analysis (PCA) and t-distributed stochastic neighbor embedding (t-SNE) were performed for visualization using the irlba package (v2.3.5.1) and Rtsne (v0.17), respectively. For the analysis of microarray-based gene expression data, batch-adjusted raw expression values were obtained from the MERAV database (http://merav.wi.mit.edu). Tumor grades were retrieved from TCGA clinical metadata, and substage labels (e.g., “IIIA”, “IIIB”) were consolidated into major stage groups (e.g., “III”) for clarity. Gene-level thresholded copy number variation (CNV) data for all TCGA tumor types were downloaded from the UCSC Xena Browser (https://xena.ucsc.edu/). CNV calls in this dataset were based on the GISTIC2 algorithm and categorized into five discrete levels (−2: deep deletion, −1: shallow deletion, 0: diploid, 1: low-level gain, 2: high-level amplification). CNV scores were integrated with the clinical and expression data using custom R scripts. All custom analysis scripts are publicly available at https://github.com/Toshi-Shigeoka/Zbtb38_paper/.

### Quantification and statistical analysis

Statistical analyses for qPCR and immunoblotting were conducted using GraphPad Prism 10 and Microsoft Excel. Unless otherwise indicated, data are presented as mean ± standard deviation (SD) from three independent experiments. Comparisons between two groups were performed using unpaired two-tailed Student’s t-tests, and one-way ANOVA was used for comparisons among more than two groups. To minimize potential biases, animals were housed in randomized cage positions, and samples were processed in a non-sequential order. Statistical significance was defined as *\*p* < 0.05, *\*\*p* < 0.01, and *\*\*\*p* < 0.001; “ns” indicates no significance (*p* > 0.05).

## Data and Code availability

All analysis scripts and processed data used for statistical analysis and figure generation are publicly available in the following GitHub repository: https://github.com/Toshi-Shigeoka/Zbtb38_paper/

## ACKNOWLEDGMENT

We thank Drs. Chio Oka and Masashi Kawaichi for their valuable suggestions and discussions. We also acknowledge Mr. Shunya Hibia, Mr. Ibuki Okuda, Ms. Fei Xu, and Ms. Tomomi Kotoku for their technical assistance. We are grateful to the members of the Ishida laboratory for their helpful suggestions and advice. This work was partly supported by the NAIST Life Science Collaboration Center (LiSCo) was supported by a Grant-in-Aid for Scientific Research (16K08587) from the Japan Society for the Promotion of Science (JSPS).

## AUTHOR CONTRIBUTIONS

T.S. planned the study, conducted the bioinformatic analysis, prepared the figures, analyzed the data, and provided technical guidance and advice. H.N. and S.T. conceived, designed, performed the experiments, and analyzed the data. E.M. conceived, designed, performed the experiments, analyzed the data, prepared the figures, and drafted the manuscript. N.N., Y.B., and Y.I. supervised the study and approved the final manuscript.

## DATA AVAILABILITY

The data used to support the findings of this study are available from the corresponding author upon reasonable request.

## RERERENCE

[1] Elmore S. Apoptosis: A Review of Programmed Cell Death. Toxicol Pathol 35, 495–516 2007.

[2] Nagata S. Apoptosis and Clearance of Apoptotic Cells. Annu Rev Immunol 36, 489–517 2018.

[3] Bock FJ & Tait SWG. Mitochondria as multifaceted regulators of cell death. Nat Rev Mol Cell Biol 21, 85–100 2020.

[4] Lossi L. The concept of intrinsic versus extrinsic apoptosis. Biochemical Journal 479, 357–384 2022.

[5] Larsen BD & Sørensen CS. The caspase-activated DNase: apoptosis and beyond. FEBS J 284, 1160–1170 2017.

[6] Sahoo G, Samal D, Khandayataray P & Murthy MK. A Review on Caspases: Key Regulators of Biological Activities and Apoptosis. Mol Neurobiol 60, 5805–5837 2023.

[7] Soldani C, Lazzè MC, Bottone MG, Tognon G, Biggiogera M, Pellicciari CE, et al. Poly(ADP-ribose) polymerase cleavage during apoptosis: When and where? Exp Cell Res 269, 193–201 2001.

[8] Czabotar PE & Garcia-Saez AJ. Mechanisms of BCL-2 family proteins in mitochondrial apoptosis. Nat Rev Mol Cell Biol 24, 732–748 2023.

[9] Siddiqui WA, Ahad A & Ahsan H. The mystery of BCL2 family: Bcl-2 proteins and apoptosis: an update. Arch Toxicol 89, 289–317 2015.

[10] Kale J, Osterlund EJ & Andrews DW. BCL-2 family proteins: Changing partners in the dance towards death. Cell Death Differ 25, 65–80 2018.

[11] Hao Q, Chen J, Lu H & Zhou X. The ARTS of p53-dependent mitochondrial apoptosis. J Mol Cell Biol 14, mjac074 (2022).

[12] Aubrey BJ, Kelly GL, Janic A, Herold MJ & Strasser A. How does p53 induce apoptosis and how does this relate to p53-mediated tumour suppression? Cell Death Differ 25, 104–113 2018.

[13] Liston P, Young SS, Mackenzie AE & Korneluk RG. Life and death decisions: The role of the IAPs in modulating programmed cell death. Apoptosis 2, 423–441 1997.

[14] Deveraux QL & Reed JC. IAP family proteins - Suppressors of apoptosis. Genes Dev 13, 239–252 1999.

[15] Dohi T, Okada K, Xia F, Wilford CE, Samuel T, Welsh K, et al. An IAP-IAP complex inhibits apoptosis. J Biol Chem 279, 34087–34090 2004.

[16] Dubrez-Daloz L, Dupoux A & Cartier J. IAPs: More than just inhibitors of apoptosis proteins. Cell Cycle 7 (8), 1036–1046 2008.

[17] Harlin H, Reffey SB, Duckett CS, Lindsten T & Thompson CB. Characterization of XIAP-Deficient Mice. Mol Cell Biol 21, 3604–3608 2001.

[18] Moulin M, Anderton H, Voss AK, Thomas T, Wong WWL, Bankovacki A, et al. IAPs limit activation of RIP kinases by TNF receptor 1 during development. EMBO J 31, 1679–1691 2012.

[19] Heard KN, Bertrand MJ & Barker PA. cIAP2 supports viability of mice lacking cIAP1 and XIAP. EMBO J 34, 2393–2395 2015.

[20] Dynek JN & Vucic D. Antagonists of IAP proteins as cancer therapeutics. Cancer Lett 332 (2), 206–214 2013.

[21] Verhagen AM, Silke J, Ekert PG, Pakusch M, Kaufmann H, Connolly LM, et al. HtrA2 promotes cell death through its serine protease activity and its ability to antagonize inhibitor of apoptosis proteins. J Biol Chem 277, 445–454 2002.

[22] Sasai N, Matsuda E, Sarashina E, Ishida Y & Kawaichi M. Identification of a novel BTB-zinc finger transcriptional repressor, CIBZ, that interacts with CtBP corepressor. Genes to Cells 10, 871–885 2005.

[23] Filion GJP, Zhenilo S, Salozhin S, Yamada D, Prokhortchouk E & Defossez P-A. A Family of Human Zinc Finger Proteins That Bind Methylated DNA and Repress Transcription. Mol Cell Biol 26, 169–181 2006.

[24] Sasai N, Nakao M & Defossez PA. Sequence-specific recognition of methylated DNA by human zinc-finger proteins. Nucleic Acids Res 38, 5015–5022 2010.

[25] Oikawa Y, Omori R, Nishii T, Ishida Y, Kawaichi M & Matsuda E. The methyl-CpG-binding protein CIBZ suppresses myogenic differentiation by directly inhibiting myogenin expression. Cell Res 21, 1578–1590 2011.

[26] de Dieuleveult M & Miotto B. DNA Methylation and Chromatin: Role(s) of Methyl-CpG-Binding Protein ZBTB38. Epigenet Insights 11, 2516865718811117 (2018).

[27] Nishii T, Oikawa Y, Ishida Y, Kawaichi M & Matsuda E. CtBP-interacting BTB zinc finger protein (CIBZ) promotes proliferation and G1/S transition in embryonic stem cells via Nanog. J Biol Chem 287, 12417–12424 2012.

[28] Kotoku T, Kosaka K, Nishio M, Ishida Y, Kawaichi M & Matsuda E. CIBZ Regulates Mesodermal and Cardiac Differentiation of by Suppressing T and Mesp1 Expression in Mouse Embryonic Stem Cells. Sci Rep 6, 34188 (2016).

[29] Oikawa Y, Matsuda E, Nishii T, Ishida Y & Kawaichi M. Down-regulation of CIBZ, a novel substrate of caspase-3, induces apoptosis. J Biol Chem 283, 14242–14247 2008.

[30] Cai Y, Li J, Zhang Z, Chen J, Zhu Y, Li R, et al. Zbtb38 is a novel target for spinal cord injury. Oncotarget 8, 45356–45366 (2017).

[31] Nishio M, Matsuura T, Hibi S, Ohta S, Oka C, Sasai N, et al. Heterozygous loss of Zbtb38 leads to early embryonic lethality via the suppression of Nanog and Sox2 expression. Cell Prolif 55, e13215 (2022).

[32] Kiefer H, Chatail-Hermitte F, Ravassard P, Bayard E, Brunet I & Mallet J. ZENON, a Novel POZ Kruppel-Like DNA Binding Protein Associated with Differentiation and/or Survival of Late Postmitotic Neurons. Mol Cell Biol 25, 1713–1729 2005.

[33] McGill GG, Horstmann M, Widlund HR, Du J, Motyckova G, Nishimura EK, et al. Bcl2 regulation by the melanocyte master regulator Mitf modulates lineage survival and melanoma cell viability. Cell 109, 707–718 2002.

[34] Pan Y, van der Watt PJ & Kay SA. E-box binding transcription factors in cancer. Front Oncol 13, 1223208 (2023).

[35] Buck-Koehntop BA, Martinez-Yamout MA, Jane Dyson H & Wright PE. Kaiso uses all three zinc fingers and adjacent sequence motifs for high affinity binding to sequence-specific and methyl-CpG DNA targets. FEBS Lett 586, 734–739 2012.

[36] Buck-Koehntop BA & Defossez PA. On how mammalian transcription factors recognize methylated DNA. Epigenetics 8, 131–137 2013.

[37] Gudbjartsson DF, Walters GB, Thorleifsson G, Stefansson H, Halldorsson BV, Zusmanovich P, et al. Many sequence variants affecting diversity of adult human height. Nat Genet 40, 609–615 2008.

[38] Kote-Jarai Z, Olama AA Al, Giles GG, Severi G, Schleutker J, Weischer M, et al. Seven prostate cancer susceptibility loci identified by a multi-stage genome-wide association study. Nat Genet 43, 785–791 2011.

[39] Jing J, Liu J, Wang Y, Zhang M, Yang L, Shi F, et al. The role of ZBTB38 in promoting migration and invasive growth of bladder cancer cells. Oncol Rep 41, 1980–1990 2019.

[40] Chen J, Xing C, Yan L, Wang Y, Wang H, Zhang Z, et al. Transcriptome profiling reveals the role of ZBTB38 knock-down in human neuroblastoma. PeerJ 7, e6352 (2019).

[41] Mead S, Uphill J, Beck J, Poulter M, Campbell T, Lowe J, et al. Genome-wide association study in multiple human prion diseases suggests genetic risk factors additional to PRNP. Hum Mol Genet 21, 1897–1906 2012.

[42] Gao X, Zhang L, Wei Y, Yang Y, Li J, Wu H, et al. Prognostic value of XIAP level in patients with various cancers: A systematic review and meta-analysis. J Cancer 10, 1528–1537 2019.

[43] Holcik M, Gibson H & Korneluk RG. XIAP: Apoptotic brake and promising therapeutic target. Apoptosis 6, 253–261 2001.

[44] De Dieuleveult M, Marchal C, Jouinot A, Letessier A & Miotto B. Molecular and clinical relevance of zbtb38 expression levels in prostate cancer. Cancers 12, 1106 (2020).

[45] Jing J, Liu J, Wang Y, Zhang M, Yang L, Shi F, et al. The role of ZBTB38 in promoting migration and invasive growth of bladder cancer cells. Oncol Rep 41, 1980–1990 2019.

