## Supplemental Figure S1-S9, supplemental Table for "Zbtb38 transcriptionally activates XIAP to regulate apoptosis in development and cancer"

### Supplementary Fig. S1

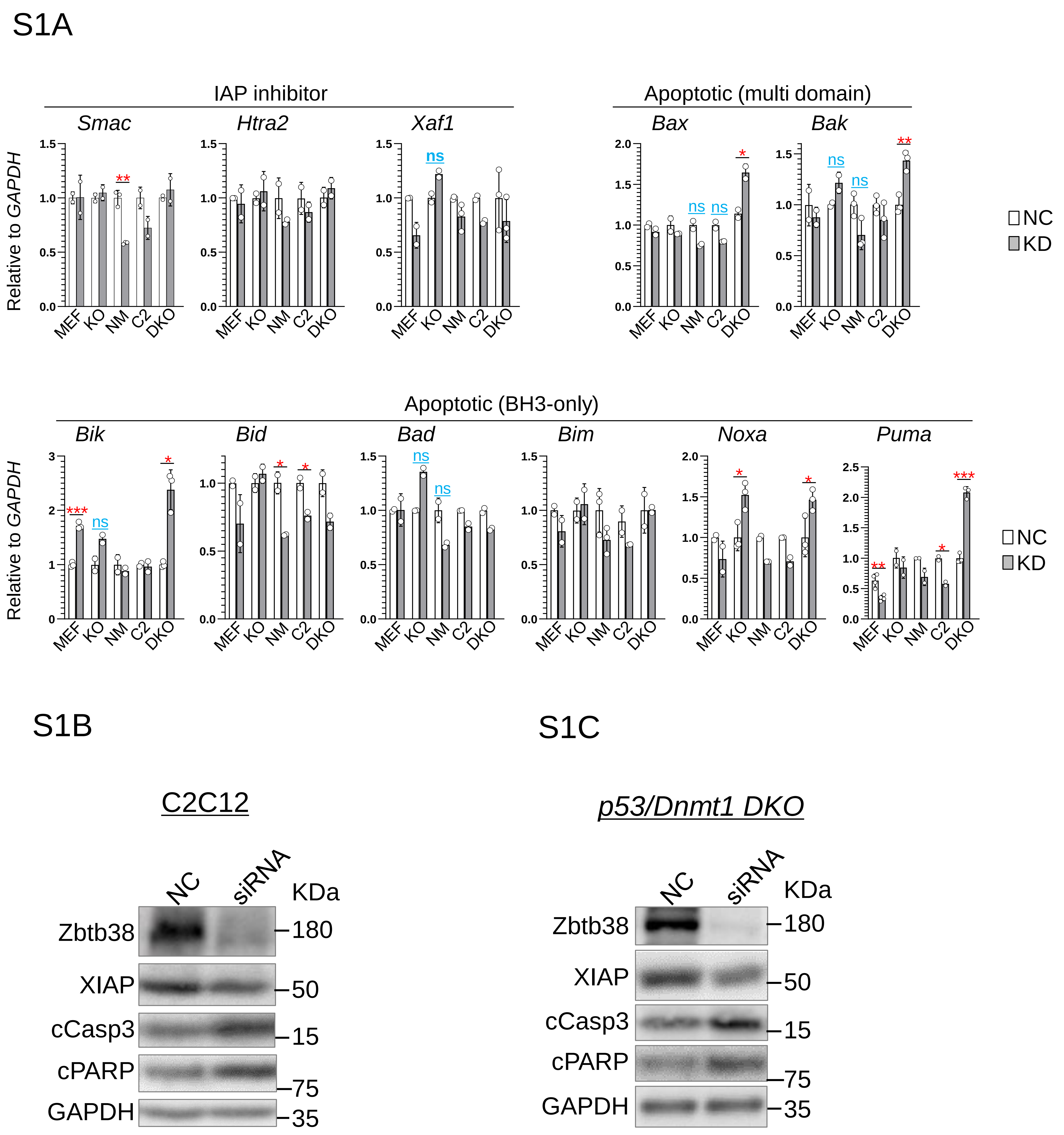


**Fig. S1 Expression of apoptosis-related genes following Zbtb38 knockdown in p53-expressing (MEFs, NMuMG, C2C12) and p53-deficient (p53 KO, DKO) cell lines.**
**S1A** mRNA levels were quantified by RT-qPCR in negative control (NC, empty bars) and Zbtb38 knockdown (KD, gray bars) groups, normalized to GAPDH (set to "1"). Abbreviations: KO, p53 knockout MEFs; NM, NMuMG; C2, C2C12; DKO, p53/Dnmt1 double-knockout MEFs. $*p < 0.05$, $**p < 0.01$, $***p < 0.001$; ns – not significant. **S1B–S1C** Zbtb38 was knockdown using siRNA in C2C12 cells (**S1B**) and p53/Dnmt1 DKO MEFs (**S1C**), and protein expression was analyzed by immunoblotting 30–36 h post-transfection. NC – negative control; siRNA – Zbtb38 siRNA duplex. GAPDH was used as a loading control.

### Supplementary Fig. S2

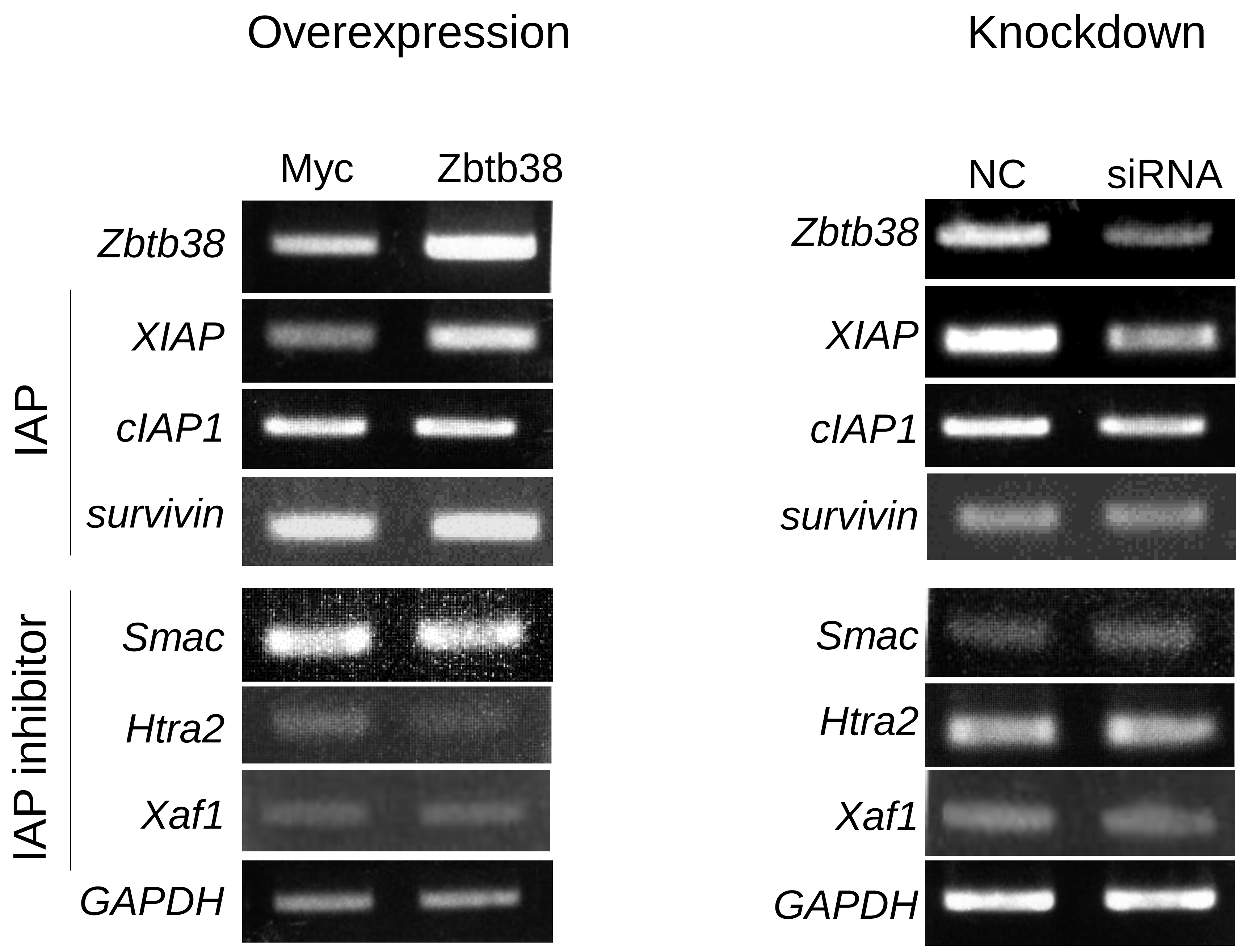


**Fig. S2 Expression analysis of apoptosis-related genes after Zbtb38 overexpression (left panel) or knockdown (right panel) in C2C12 cells.**
Cells were transfected with Myc or Myc-Zbtb38 for 48 h (left panel), or with a negative control (NC) or siRNA against Zbtb38 for 36 h (right panel). Semi-quantitative RT-PCR was used to analyze the mRNA levels of the indicated genes. GAPDH was used as an internal control.

Supplementary Fig. S3

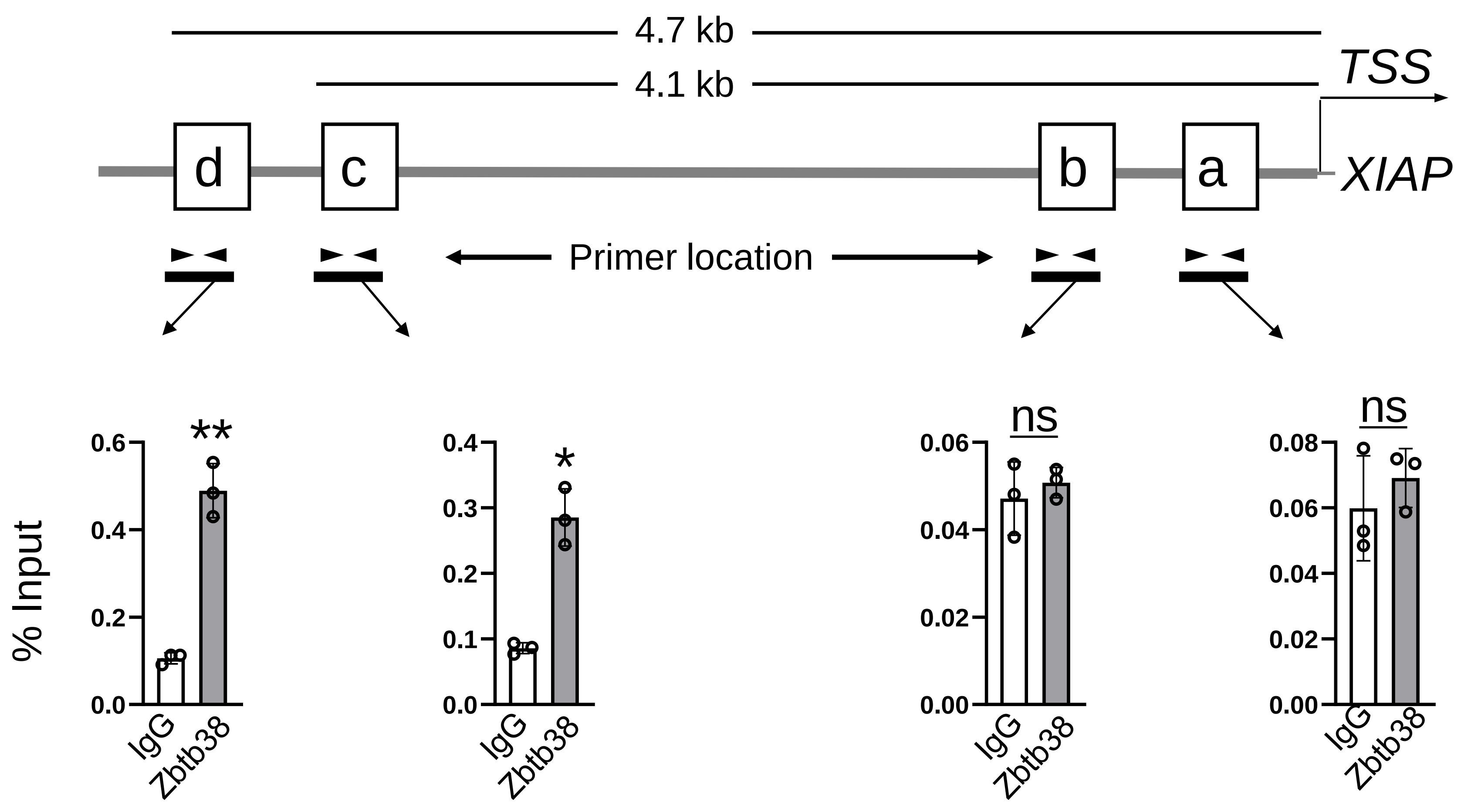


**Fig. S3 ChIP-qPCR results for *p53* KO MEFs.**
Schematic representation of the regulatory regions of *XIAP*. Regions a and b represent putative promoter regions, and regions c and d indicate the enhancer regions of XIAP. TSS, transcriptional start site. IgG-precipitated DNA (negative control) and Zbtb38-immunoprecipitated DNA were amplified by qRT-PCR using primers at the indicated locations. Data are shown as fold enrichment relative to the input control, which was set to 1. *$p < 0.05$, **$p < 0.01$. "ns" indicates no significance.

Supplementary Fig. S4

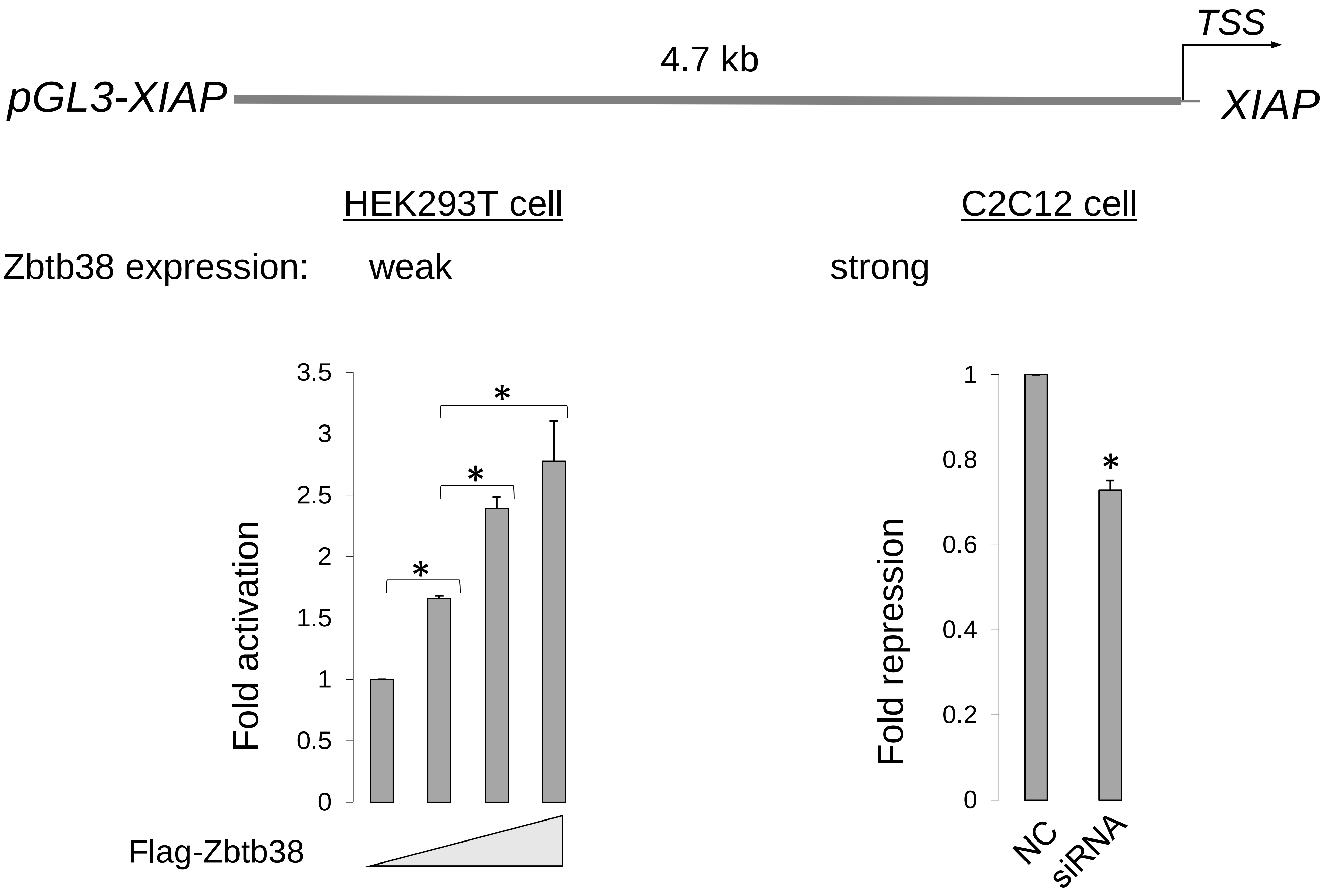


**Fig. S4 Zbtb38 overexpression in HEK293T cells and knockdown in C2C12 cells.** Top panel: schematic of the pGL3-XIAP. Left panel: pGL3 and pGL3-XIAP were co-transfected with expression vectors for Flag or increasing amounts of Flag-Zbtb38 into HEK293T cells for 40-48 h. Luciferase activity was determined relative to that of Flag/pGL3, which was set to 1. Right panel: pGL3 vector and pGL3-XIAP were co-transfected with a negative control (NC) or siRNA against Zbtb38 for 48 h. Luciferase activity was determined relative to that of pGL3, which was set to 1. *$p < 0.05$.

Supplementary Fig. S5

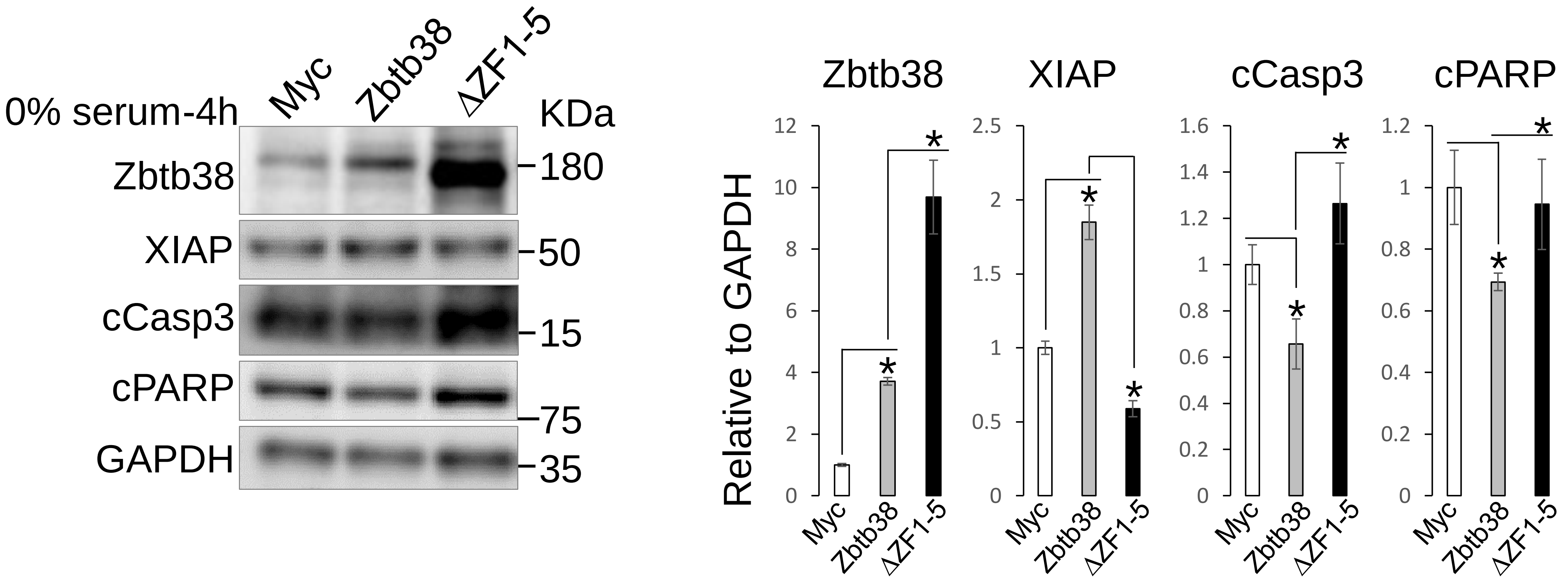


**Fig. S5 ZF1-5 of Zbtb38 is required for inhibiting apoptosis in C2C12 cells.**
Myc, Myc-Zbtb38, and Myc-ΔZF1-5 were transfected into C2C12 cells for 36 h and then switched to medium without FBS for 4 h. Data were quantified relative to GAPDH, which was set to 1, and are shown in the right panel. *$p < 0.05$.

Supplementary Fig. S6

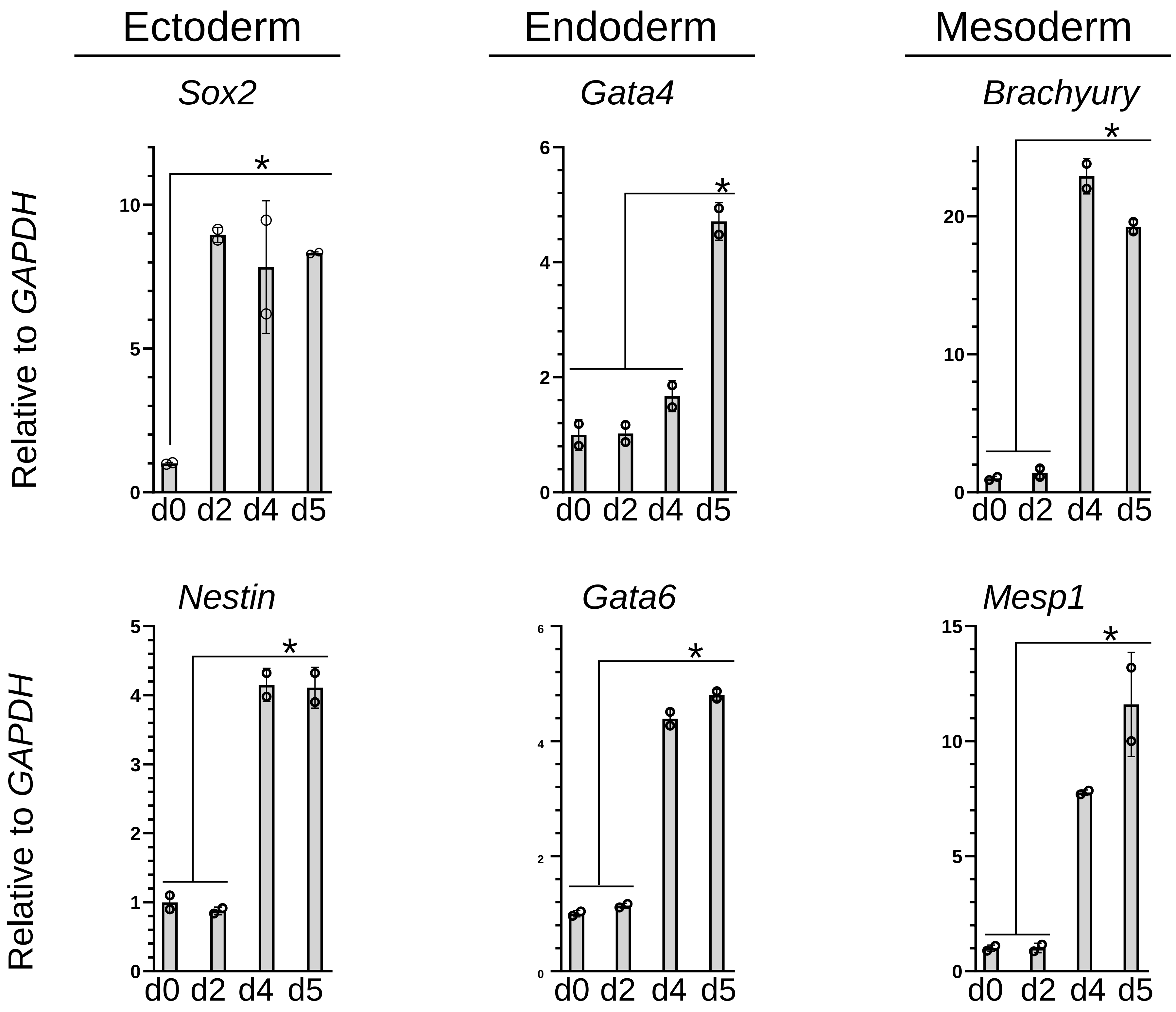


**Fig. S6 RT-qPCR results of the indicated gene expression in wild-type ES cells at the indicated differentiation days.**
ES cells were cultured in ES medium to maintain an undifferentiated state (d0) and then differentiated into embryoid bodies in suspension culture lacking leukemia inhibitory factor. The data are representative of three independent experiments. Transcript levels were normalized to those of GAPDH. *$p < 0.05$.

Supplementary Fig. S7

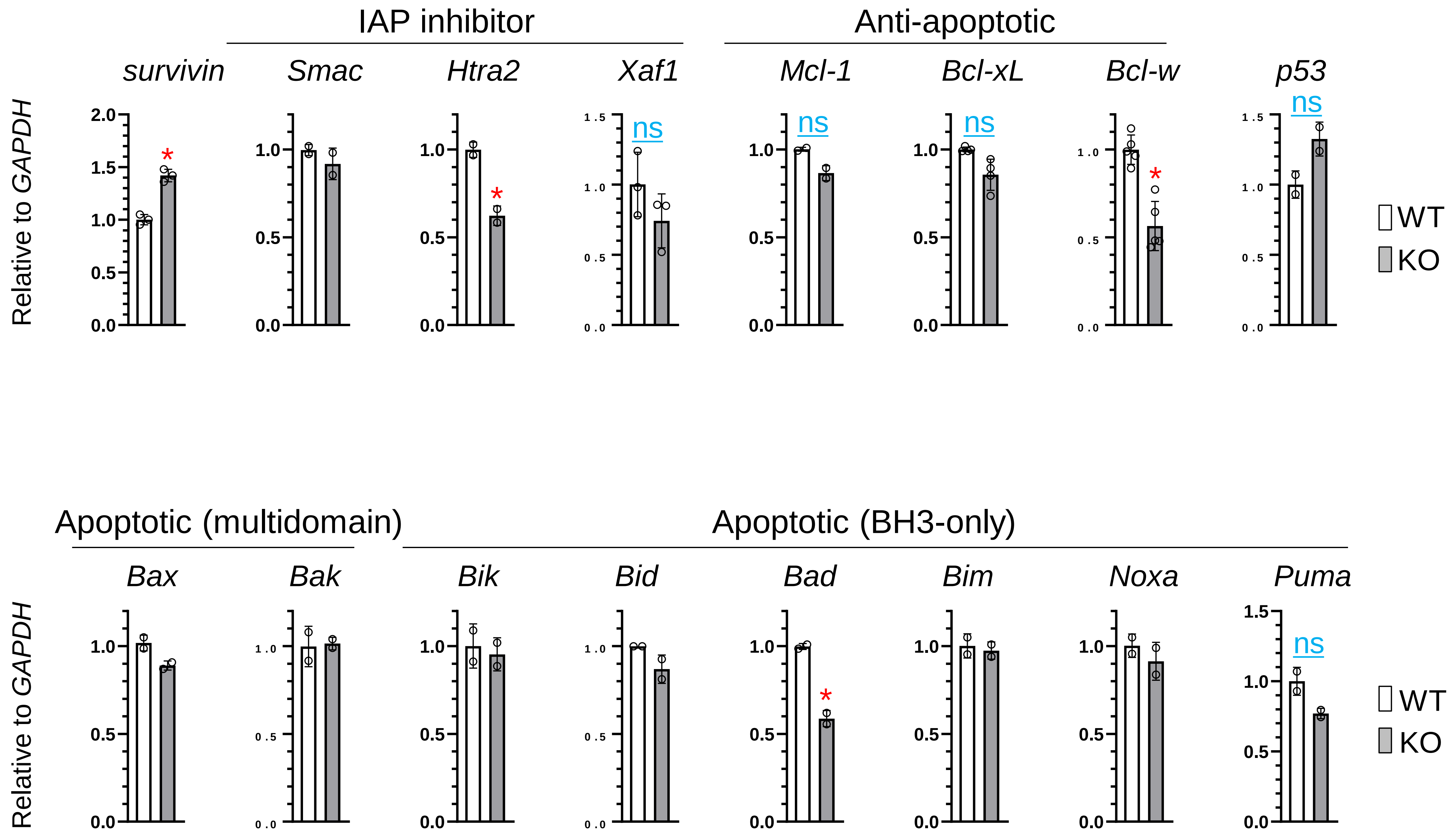


**Fig. S7 RT-qPCR analysis of the expression of indicated genes in undifferentiated ES cells.** WT (empty column) and KO (Zbtb38 knockout, gray column) ES cells were used. Each transcript was normalized to that of GAPDH, which was set to 1. *$p < 0.05$. "ns" indicates no significance.

Supplementary Fig. S8

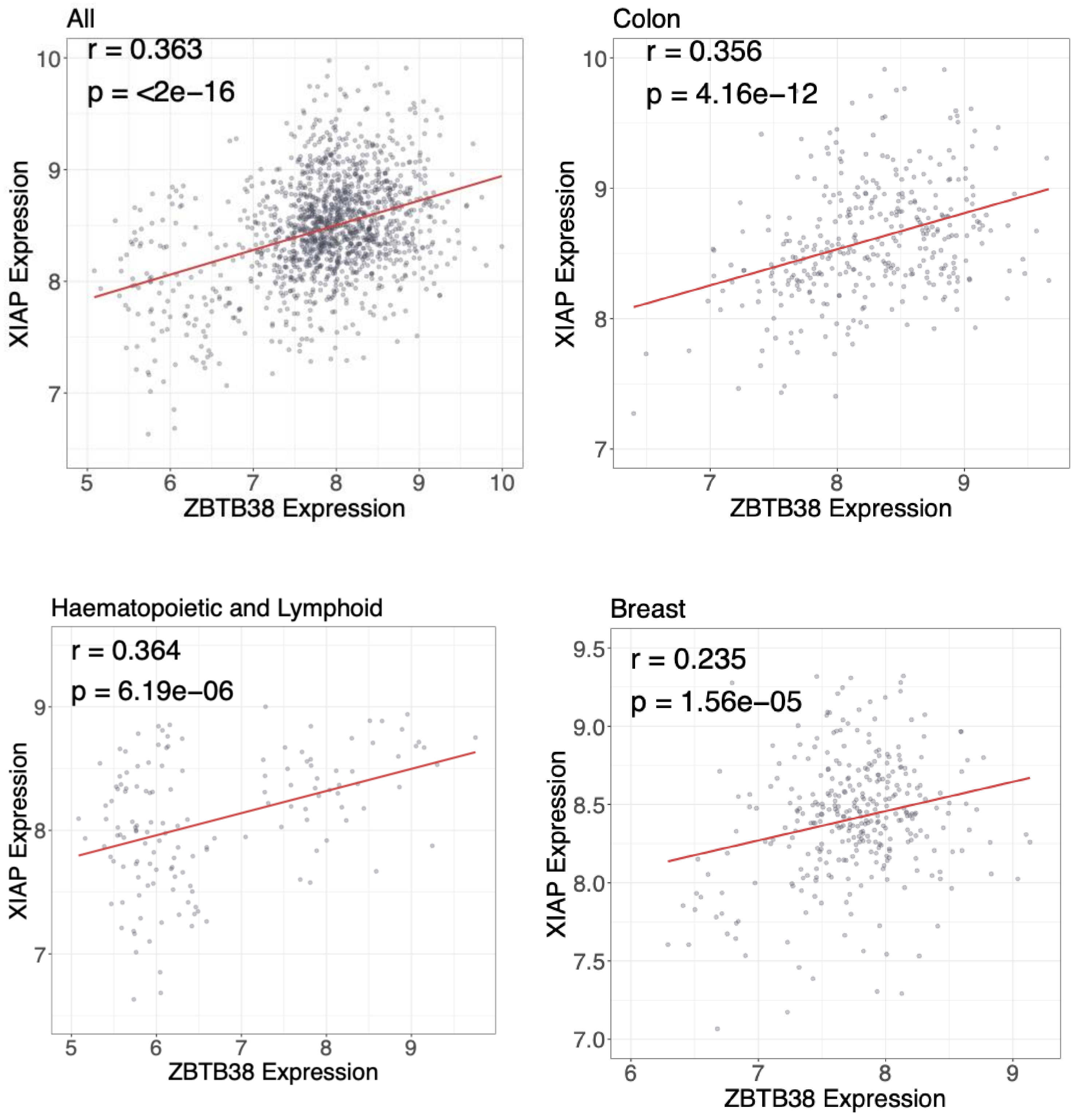


**Fig. S8 Expression analysis of ZBTB38 in human tumors based on MERAV.** Scatter plots showing the correlation between ZBTB38 and XIAP expression across all TCGA tumor types (**A**), in colon tumors (**B**), and in haematopoietic and lymphoid tumors (**C**) and breast tumors (**D**). Pearson's correlation coefficient (r) and p-values are indicated in each plot.

### Supplementary Fig. S9

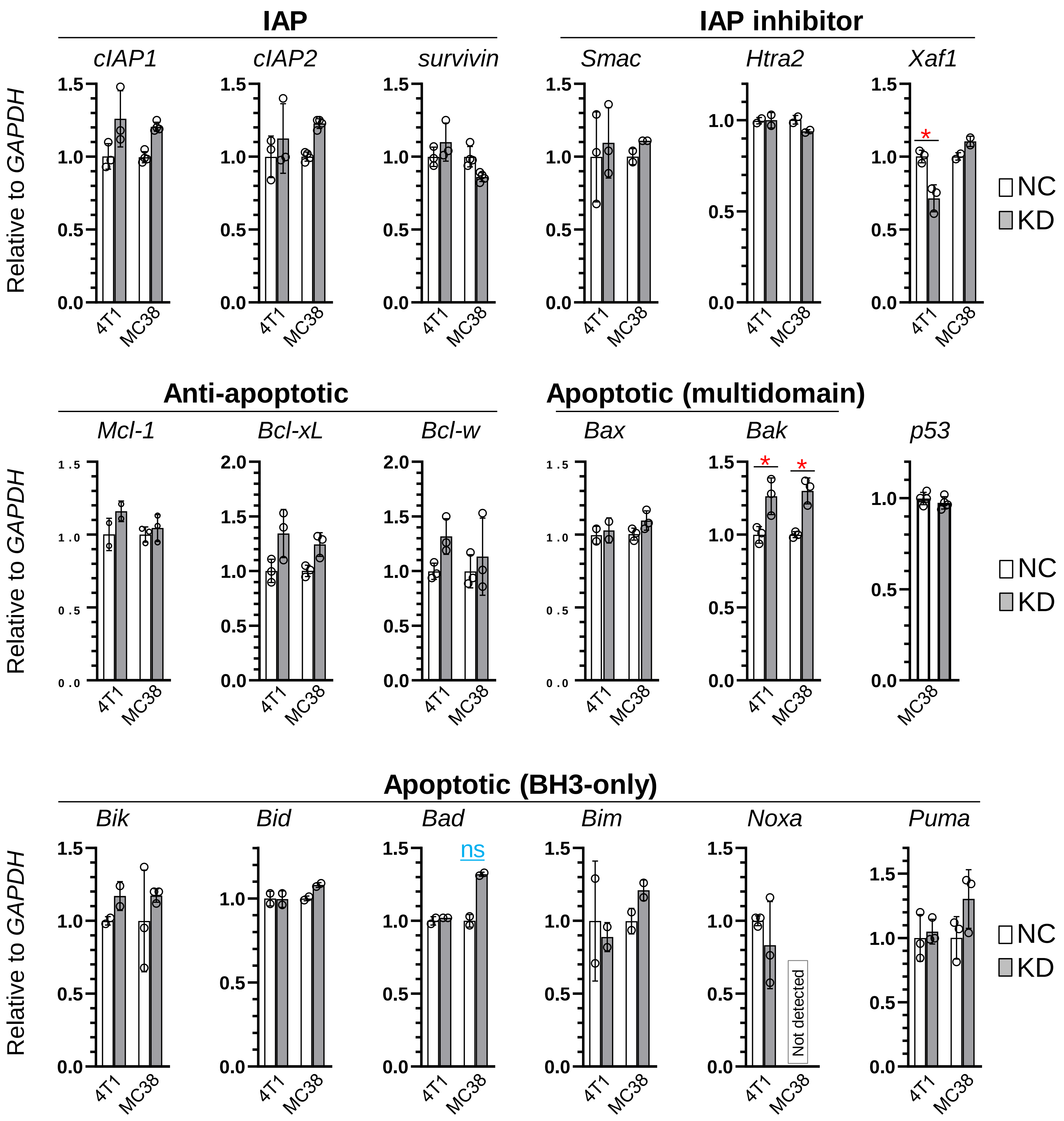


**Fig. S9 RT-qPCR analysis of gene expression in 4T1 and MC38 cells.** WT (empty column) and KO (Zbtb38 knockout, gray column) cells were used. Each transcript was normalized to GAPDH, set to 1. $p < 0.05$. "ns" indicates no significance.

**Supplemental table**

**Primer sequences for RT-qPCR**

| Gene | Primer sequence (5′ to 3′) | Product size |
|---|---|---|
| Zbtb38 | F: ACGCTCAAGATCCACGAGAG<br>R: ACCCTGAAAGCAGAACTGACA | 69 bp |
| XIAP | F: AACAAAGTGGGAAGGTGAGAG<br>R: TGTGTATGTGTCTGGGTGTAAG | 101 bp |
| cIAP1 | F: ATTGGAAGAGCAGTTGCGG<br>R: CAGTCCCCTTGATTGTCCCC | 266 bp |
| cIAP2 | F: CTGGCTATTTCAGTGGCTCTTA<br>R: TGCAAAGTGGTAGGGACTTG | 105 bp |
| Bcl-2 | F: CTTCGCAGAGATGTCCAGTC<br>R: AGGGCGATGTTGTCCACCAG | 192 bp |
| Bax | F: ATGCGTCCACCAAGAAGCTGAG<br>R: CCCCAGTTGAAGTTGCCATCAG | 166 bp |
| Bcl-xL | F: GTGAAGCAAGCGCTGAGAG<br>R: AACTTGCAATCCGACTCACC | 250 bp |
| Noxa | F: AGTTCGCAGCTCAACTCAGG<br>R: GCCGTAAATTCACTTTGTCTCC | 201 bp |
| Puma | F: AGACAAGAAGAGCAGCATCG<br>R: CTAGTTGGGCTCCATTTCTGG | 117 bp |
| Bik | F: CTATACACAGACTCGCTGTCAC<br>R: AAGACTCTCCAGGACCAGAT | 113 bp |
| Bid | F: TCACAGACCTGCTGGTGTTC<br>R: TGTCTGGCAATGTTGTGGAT | 220 bp |
| Mcl-1 | F: CCTTTACTGTTGGCGTGTTATG<br>R: GGGTGAGAGTTCTAATGCAGAT | 109 bp |
| Bim | F: CTTCCATACGACAGTCTCAG<br>R: TCTTCAGCCTCGCGGTAATC | 142 bp |
| Htra2 | F: CGTGGAATACATTCAGACCG<br>R: AAACTCCCTAAGGCGATCAG | 151 bp |
| Smac | F: TTTCCTGTCTTGGCTAACTC<br>R: CTCCGATTTCTGAGCAATAGG | 123 bp |
| survivin | F: GCTGTACCTCAAGAACTACCG<br>R: GTTCCCAGCCTTCCAATTCC | 173 bp |
| p53 | F: GGGAATAGGTTGATAGTTGTCAG<br>R: GGGTGAGATTTCATTGTAGGTG | 94 bp |
| Bak | F: AGATGATATTAACCGGCGCTAC<br>R: GGCGATCTTGGTGAAGAGTT | 101 bp |
| Xaf1 | F: CTGCGCTTCATAGTCCTTTGCC<br>R: CTTGGCAGGGTGCTGTTGG | 117 bp |
| Bcl-w | F: CAAGTGCAGGATTGGATGGTGG<br>R: CTGTCCTCACTGATGCCCAGTT | 157 bp |
| Bad | F: AGACGCTAGTGCTACAGATAGG<br>R: CGTCCCTGCTGATGAATGTT | 202 bp |
| GAPDH | F:CAATGTGTCCGTCGTGGATCT<br>R:GTCCTCAGTGTAGCCCAAGATG | 124 bp |

**Primers used for semi-quantitative PCR**

| Gene | Primer sequence (5′to 3′) | Product size |
|---|---|---|
| XIAP | F: TCACTTGAGGTCCTGATTGC<br>R: GCTTGAACGTAATGACGGTG | 305 bp |
| cIAP1 | F: GCACAGACAGTTCTATCCCA<br>R: GCTCTGACATAGCATCATCC | 351 bp |
| survivin | F: GCTGTACCTCAAGAACTACCG<br>R: GTTCCCAGCCTTCCAATTCC | 167 bp |
| Smac | F: TTTCCTGTCTTGGCTAACTC<br>R: CTCCGATTTCTGAGCAATAGG | 210 bp |
| Htra2 | F: CGTGGAATACATTCAGACCG<br>R: AAACTCCCTAAGGCGATCAG | 298 bp |
| Xaf1 | F: CTTCATAGTCCTTTGCCCAG<br>R: GATCTTCGATTTCCAGAGTCC | 317 bp |
| Zbtb38 | F: CCATAGATCACAGACTCTCCAT<br>R: CTGTAGCTGATCACAGAGGCCGAG | 511 bp |
| GAPDH | F: CCATCACCATCTTCCAGGAG<br>R: CCTGCTTCACCACCTTCTTG | 577 bp |

**Primers used for ChIP assay**

| location | Primer sequence (5′to 3′) |
|---|---|
| **a** | F: GAACTTTGCCTTGAATATGTAATG<br>R: CAGAGCAGAATTACTGATC |
| **b** | F: CAAACAGATTGTTCCTTGACCAACAG<br>R: TGATGGAAGTGGACCATTCCT |
| **c** | F: CTATCATTATCCAGCTTATGCTGATTC<br>R: CACATATCTCTGAAGAGCTGAC |
| **d** | F: GGACGTAATCTTCATAATGTA<br>R: GTCACATAAAGGCGAACTTCC |

**Primers used for the generation of reporter constructs**

pGL3-XIAP 4.7 kb
F: CGGGGTACCCATTGGCCTTAATTCCACTG
R: CCGCTCGAGGGACTATTTCTGATACTAAGCAC

pGL3-XIAP m1
TGAGTGTTCCAGTGCTGGACAGTAA
TTACTGTCCAGCACTGGAACACTCA

pGL3-XIAP m1/m2
CAGAATGTTTTCAAAGTGCTGGAAAAATGACAG
GTGTCATTTTTCCAGCACTTTGAAAACATTCTG
